# Functional diagnostics using fresh uncultured lung tumor cells to guide personalized treatments

**DOI:** 10.1101/2020.08.12.247817

**Authors:** Sarang S. Talwelkar, Mikko I. Mäyränpää, Lars Søraas, Swapnil Potdar, Jie Bao, Annabrita Hemmes, Nora Linnavirta, Jon Lømo, Jari Räsänen, Aija Knuuttila, Krister Wennerberg, Emmy W. Verschuren

## Abstract

Functional profiling of a cancer patient’s tumor cells holds potential to tailor personalized cancer treatment. Here we report the utility of Fresh Uncultured Tumor-derived EpCAM+ epithelial Cells (FUTC) for *ex vivo* drug response interrogation. Analysis of murine *Kras* mutant FUTCs demonstrated pharmacological and adaptive signaling profiles comparable to subtype-matched cultured cells. Applying FUTC profiling on non-small cell lung cancer patient samples, we generated robust drug response data in 18 of 19 cases, where the cells exhibited targeted drug sensitivities corresponding to their oncogenic drivers. In one of these cases, an *EGFR* mutant lung adenocarcinoma patient refractory to osimertinib, FUTC profiling was used to guide compassionate treatment. FUTC profiling identified selective sensitivity to disulfiram and the combination of carboplatin plus etoposide and the patient received substantial clinical benefit from the treatment with these agents. We conclude that FUTC profiling provides a robust, rapid, and actionable assessment of personalized cancer treatment options.

## INTRODUCTION

The concept of precision medicine means giving the right drug to the right individual at the right time. In this context, genotype-guided pairing of tumors with drugs that target tumor-selective driver mutations has been in clinical practice for more than two decades. For non-small cell lung cancer (NSCLC) patients, inhibitors targeting EGFR, ALK, BRAF, MET, and ROS are approved for genotype-guided indications that match for around 30% of the patients.^1^ However, even with the genetic matches, only 50-70% of treated patients benefit from these treatments. When attempting to use genotype-guided therapies beyond currently approved indications, trials have shown that only 10% of patients can be paired with genomics-matched targeted treatments, and at best one third of these receives clinical benefit.^2, 3^ Hence, prediction of successful precision anticancer treatments at the individual level remains challenging, largely because of extensive genomic and phenotypic heterogeneity of many cancer types, including NSCLC. Therefore, to identify precision medicines for the majority of NSCLC patients, we need additional tools beyond those that are currently used in the clinic.

Combining static genetic measurements with pharmacological interrogation of patient-derived cancer cells can provide a more comprehensive approach for predicting effective treatments; ^4–6^ however, proof of its clinical utility is lacking particularly for solid tumors. For patients with hematological malignancies, we and others have successfully implemented individualized treatment strategies guided by *ex vivo* drug-response testing of patient biopsies. ^7, 8^ Based on these early successes of *ex vivo* testing to tailor patient treatments, multiple clinical trials were initiated for leukemia patients (e.g. NCT01620216 and NCT04267081). For solid tumors, similar functional diagnostic methodologies can however not readily be adopted; reasons include the fragile and short-lived nature of tissues, for example observed in organotypic tumor slice cultures, ^9, 10^ and a general lack of robust methods to isolate and propagate primary epithelial cells.

In promising recent developments, new culture and organoid approaches enabling the long-term expansion of primary epithelial cells have been introduced. ^11, 12^ Towards translational applications, the use of conditionally reprogrammed (CR) cultures derived from clinical lung tumors successfully identified novel actionable treatments and molecular mechanisms underlying drug resistance. ^4, 13, 14^ Similarly, NSCLC organoids have been shown to recapitulate oncogenic addictions and tissue architecture of the original tumors, offering an optional model for exploring inter- and intra-tumoral functional heterogeneity. ^15, 16^

While the implementation of CR or organoid cultures in principle permits drug response profiling of patient-derived cells, these models take long times to be established, and importantly, do not guarantee the expansion of malignant cells. ^4, 15, 17^ Their functional interrogation will therefore unlikely lead to significant impacts on personalized diagnostics and patient care. ^11^ To circumvent challenges associated with *ex vivo* cultures, we tested the utility of Fresh Uncultured Tumor-derived Cells (FUTCs) for drug response assessment. Using this approach, we present a diagnostic assay for the rapid identification of actionable treatments using patient-derived tumor cells.

## RESULTS

### Tumor-derived epithelial cultures often do not represent malignant cells

With the aim to identify individualized treatment options, and to understand genetic driver - drug response relationships, we established NSCLC patient-derived CR cultures to allow for pharmacogenomic screening. To confirm their identity, we performed targeted next-generation sequencing for 578 cancer-related genes in 11 primary cultures, as well as matched tumor and adjacent normal lung tissues. Altogether, 71 nonsynonymous somatic mutations in 47 genes were detected in tumor tissues (Table 1). Nine of the 11 tested cultures lacked the oncogenic mutations detected in the reference tumor. Notably, the two cultures that showed concordance of mutations with their respective source tissue were derived from different regions from the same tumor (Table 1). The success rate of establishing patient-derived malignant cultures was therefore limited to about 10%, arguing that CR culture establishment is not an effective approach for generating pharmacogenomic screening information from NSCLC patient samples.

**Table 1.**
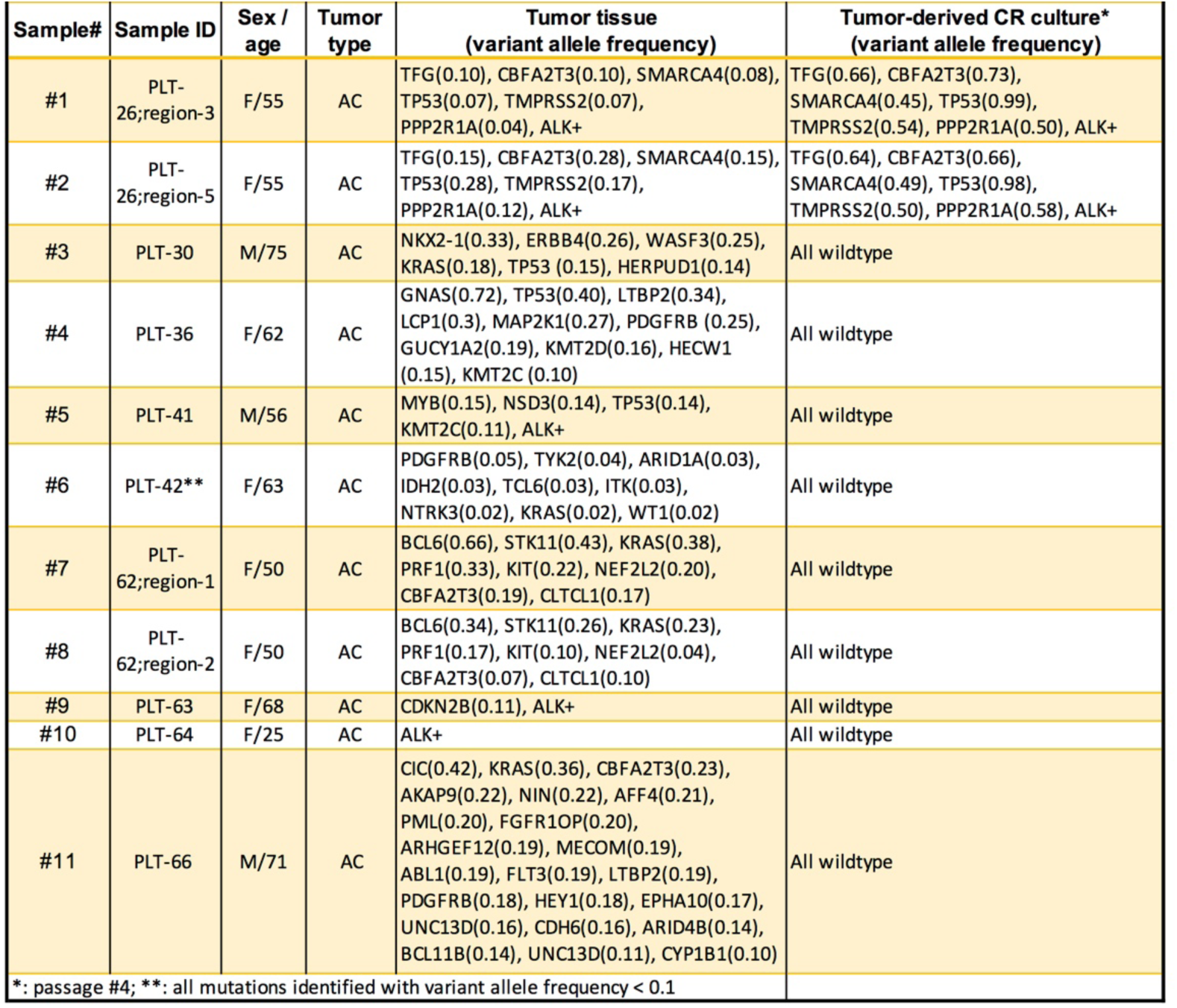
Genomic analysis of NSCLC tumor tissue and tumor-derived CR cultures

### Murine fresh uncultured tumor-derived cells (FUTCs) capture drug responses of other ***ex vivo* cell culture systems**

To address the challenge of cell culture adaptation, we investigated whether freshly isolated EpCAM-expressing epithelial cells can directly be utilized for drug response assessment. We first performed functional profiling of FUTCs and CR cultures derived from murine NSCLC tumors obtained from *Kras^G12D^;Lkb1^fl/fl^* (KL) and *Kras^G12D^;p53^fl/fl^* (KP) models. These models were selected because (i) they represent tumors with common genetic drivers and histopathological diversities detected in clinical specimens, ^18^ (ii) they allow CR culture establishment with relative ease, permitting comparisons of tumor-matched FUTCs and cultures (Fig. 1A), and (iii) we have gained understanding on their tumor subtype-selective drug sensitivities. ^19^

**Figure 1.**
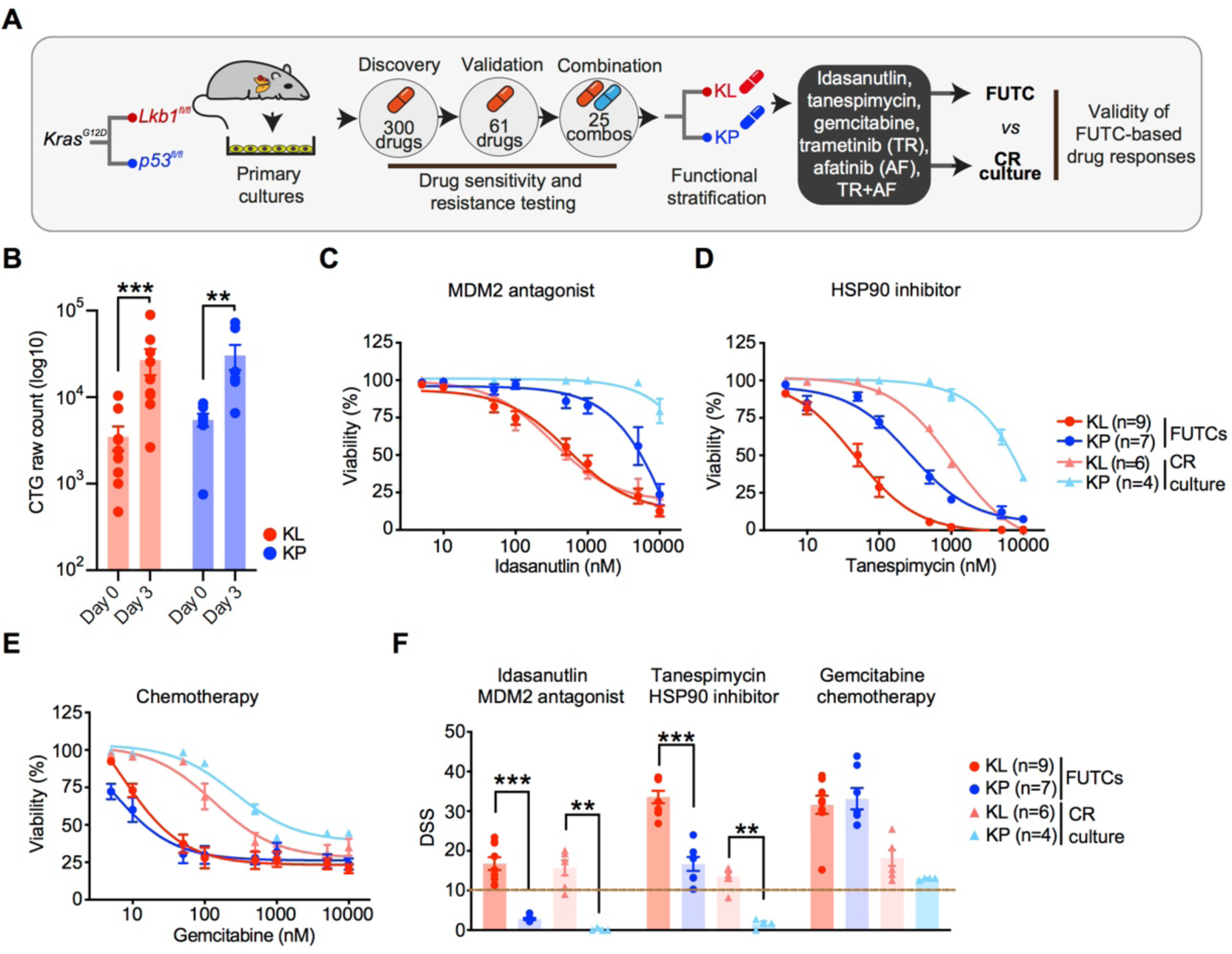
Murine FUTCs exhibit functional profiles comparable to conditionally reprogrammed cultures. (A) Schematic of comprehensive drug testing of KP and KL tumor-derived cultures to validate the ability of FUTCs to identify subtype-selective treatments. (B) Viability assessment of FUTCs at 0 and 72 h. Dose-response curves of tumor-matched FUTCs and CR cultures treated for 72 h with (C) idasanutlin (MDM2 antagonist), or (D) tanespimycin (HSP90 inhibitor), or (E) gemcitabine (chemotherapeutic). (F) DSS calculated for (C-E) and compared between KL and KP subtypes for both FUTCs and CR cultures. Data are represented as means ± SEM. Student’s *t* test *p* values are *<0.05, **p<0.01, ***p<0.001.

First, we confirmed that FUTCs survive and grow during culture by measuring cellular ATP levels using CellTiter-Glo (CTG). FUTCs from the different tumor groups exhibited significantly higher CTG readouts after three days of culture (Fig. 1B). Next, we assessed the utility of FUTCs to predict known genotype-selective drug sensitivities. We confirmed that only p53-expressing FUTCs responded to the Mdm2-p53 interaction inhibitor idasanutlin (Fig. 1C and F). Moreover, KL-derived FUTCs exhibited selective responses to HSP90 inhibition (Fig. 1D and F), corroborating findings from cultured lung cancer models. ^19–21^ Lastly, we confirmed that gemcitabine, an approved chemotherapeutic agent for NSCLC treatment, potently inhibited the viability of both KL and KP cells (Fig. 1E-F), matching previous findings in CR cultures. ^19^ In further agreement with published data, ^22, 23^ a synergistic interaction was detected between MEK and ERBB receptor family inhibition in KL, but not in KP FUTCs (Fig. 2A). We also could confirm adaptive reactivation of the MAPK and PI3K signaling pathways, detected by the re-phosphorylation of MEK and AKT after extended MEK inhibition, described in many other NSCLC studies including our murine CR cultures (Fig. 2B-D and S1).

**Figure 2.**
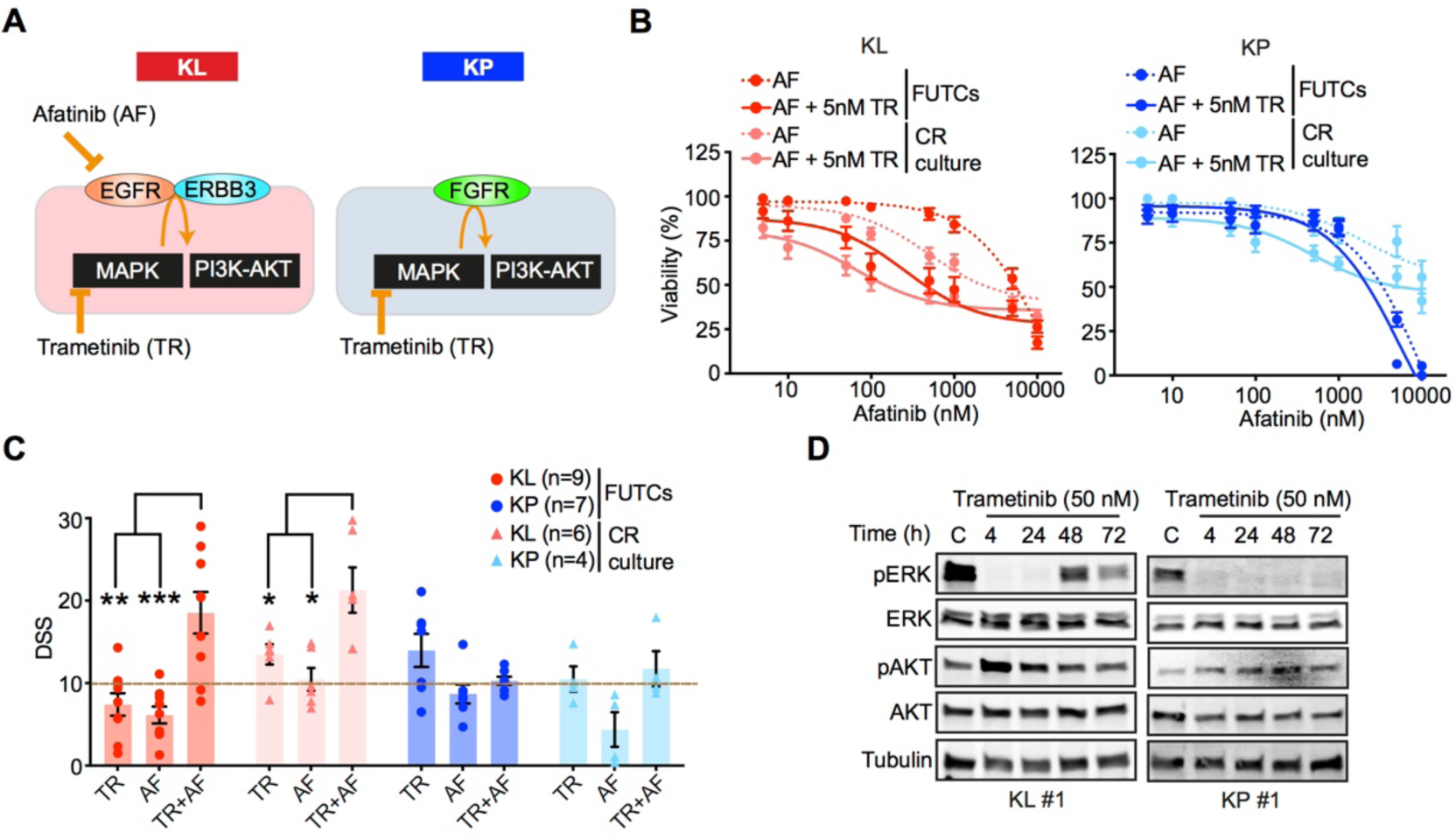
Murine FUTCs show treatment-adaptive signaling mechanisms. (A) Graphical model for subtype-selective adaptive activation of MAPK and PI3K-AKT pathways in murine *Kras* mutant NSCLC cultures. (B) Dose-response curves of FUTCs and CR cultures treated trametinib (TR), afatinib (AF), and combination treatment. For the combination screen, 5 nM of trametinib was used together with a dose series of afatinib. (C) DSS calculated for (B) and compared between KL and KP subtypes for both FUTCs and CR cultures. (D) Immunoblots of KL and KP FUTCs treated with vehicle (C; DMSO) and or treated with 50 nM trametinib for various time points (4, 24, 48, and 72 h), and probed with indicated antibodies. Error bars represent ± SEM. Student’s *t* test *p* values are *<0.05, **p<0.01, ***p<0.001.

### Patient-derived FUTCs are amenable for pharmacological profiling

To test a FUTC-based approach with patient-derived cells, we conducted a drug sensitivity and resistance testing (DSRT) study on 19 clinical NSCLC samples (Table S1). To identify drug responses selective to tumor-derived EpCAM+ cells, responses were compared to tumor-derived non-epithelial EpCAM– cells, and also to healthy lung tissue-derived EpCAM+ cells as tissue-matched controls (Fig. 3A). Similar to the murine FUTCs, the patient-derived FUTCs exhibited good viability and metabolic stability in culture (Fig 3B and S2A-S2B). Furthermore, analysis of *KRAS* mutations in six patient samples revealed an average 3.7-fold enrichment of cancer cells in tumor-derived EpCAM+ cell fractions compared to matched bulk tumor tissues (Fig. 3C). Similarly, EpCAM+ fractions derived from samples carrying an ALK rearrangement exhibited enrichment of *ALK* fusion-carrying cells (Fig. S2C).

**Figure 3.**
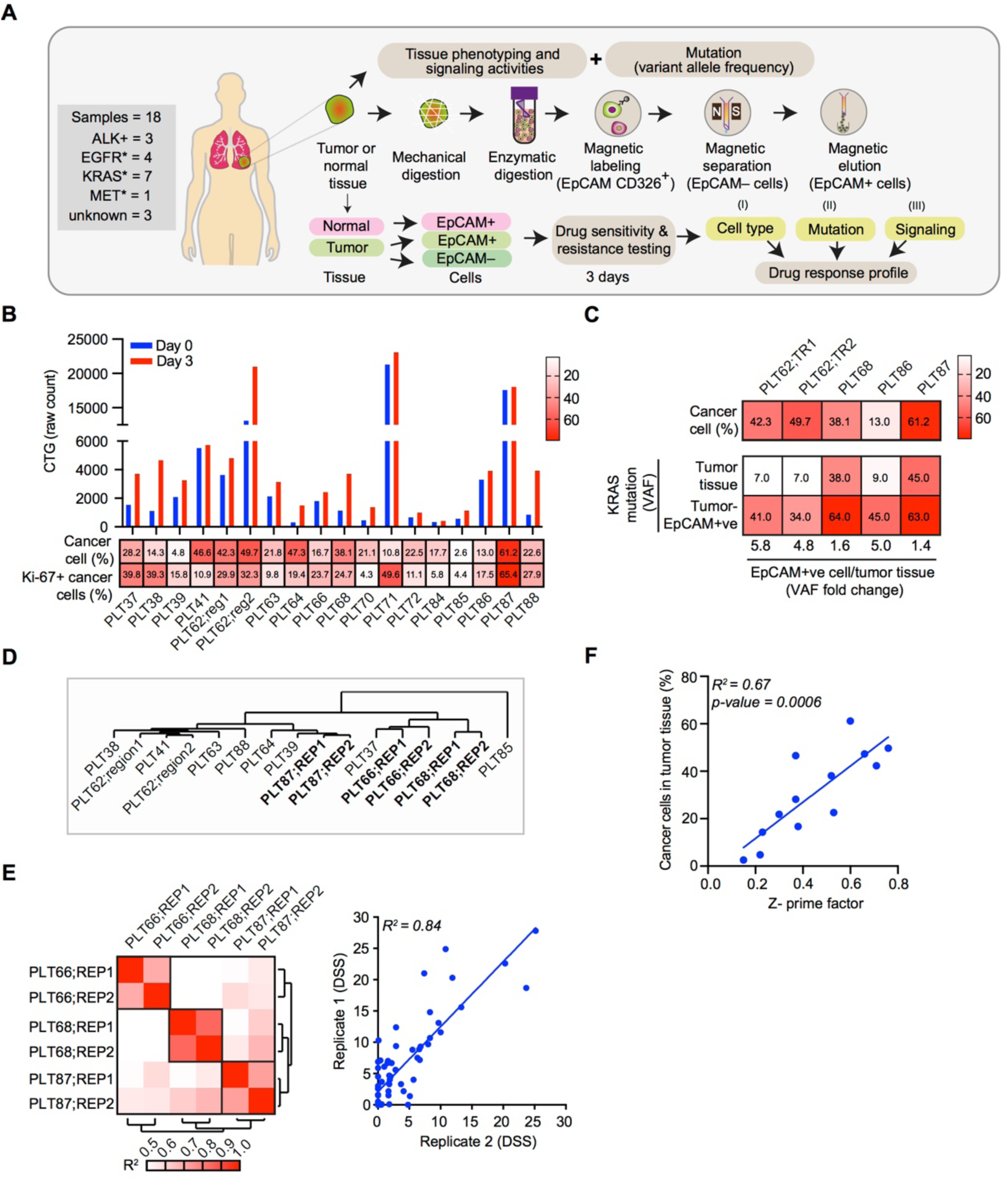
Application of patient-derived FUTCs for pharmacological screening. (A) Schematic of FUTC-based drug sensitivity and resistance testing from clinical specimens. (B) Viability assessment of patient-derived cells at days 0 and 3 after seeding in 384-well plates. Heatmap showing percentages of total cancer cells and Ki-67 positive cancer cells normalized to total cancer cells in the tumor tissue. (C) Heatmap representing *KRAS* mutation variant allele frequencies in patient-matched tumor tissues, tumor-derived EpCAM+ and EpCAM– cell populations and percentage of cancer cells. (D) DSS (66 compounds) of tumor-derived EpCAM+ cells were clustered by using a complete linkage method, coupled with Euclidean distance measurement. (E) Heatmap of Pearson’s correlation coefficient values between technical replicates of the same sample, or samples from different patients (left). Representative correlation plots of DSS values, comparing technical replicate screens of PLT68 (right). (F) Correlation plot comparing the association between the percentage of cancer cells in the tumor tissue and Z-factors obtained from respective DSRT screens.

Next, we performed screening with 66 lung cancer-selective drugs on 13 samples, or with smaller compound sets on five samples where only limited EpCAM+ FUTCs could be isolated, and for 18/19 samples we observed robust drug screening data (Z’>0.2). Unsupervised hierarchical clustering of the drug sensitivity scores revealed that technical replicates from the same patient sample cluster together, thus confirming robustness of the screening data generated using FUTCs (Fig. 3D-E). Finally, through multifactorial analyses, we showed that the robustness of the screening data correlated with the cancer cell fraction in the tumor tissue (Fig. 3F and S2D). Together, these results indicate that FUTCs are relevant *ex vivo* models of tumor tissue, and that this single-cell population can be diagnostically profiled prior to culture establishment.

### Responses to approved targeted therapies are exposed with patient-derived FUTCs

To further validate diagnostic use of the FUTC assay, responses to well-established targeted receptor tyrosine kinase inhibitors were analyzed. Tumor-derived EpCAM+, but not EpCAM–, cells from three *EGFR* mutant patient samples (L858R and E746-A750del) responded to several classes of EGFR inhibitors. Third-generation mutant-selective osimertinib selectively and most effectively reduced the viability of *EGFR* mutant EpCAM+ tumor cells (drug sensitivity score [DSS]>10), while the second-generation pan-ERBB inhibitor afatinib reduced the viability of both tumor and normal EpCAM+ cells (Fig 4A-B). These results are consistent with the known superior selectivity of osimertinib towards mutant *EGFR*. ^24, 25^ Similarly, in FUTCs derived from a clinical sample carrying an activating exon 14 skipping mutation in the *MET* gene, the ALK/MET inhibitor crizotinib showed strongest sensitivity (DSS>10), and this was selective for EpCAM+ tumor cells (Fig 4C-D).

**Figure 4.**
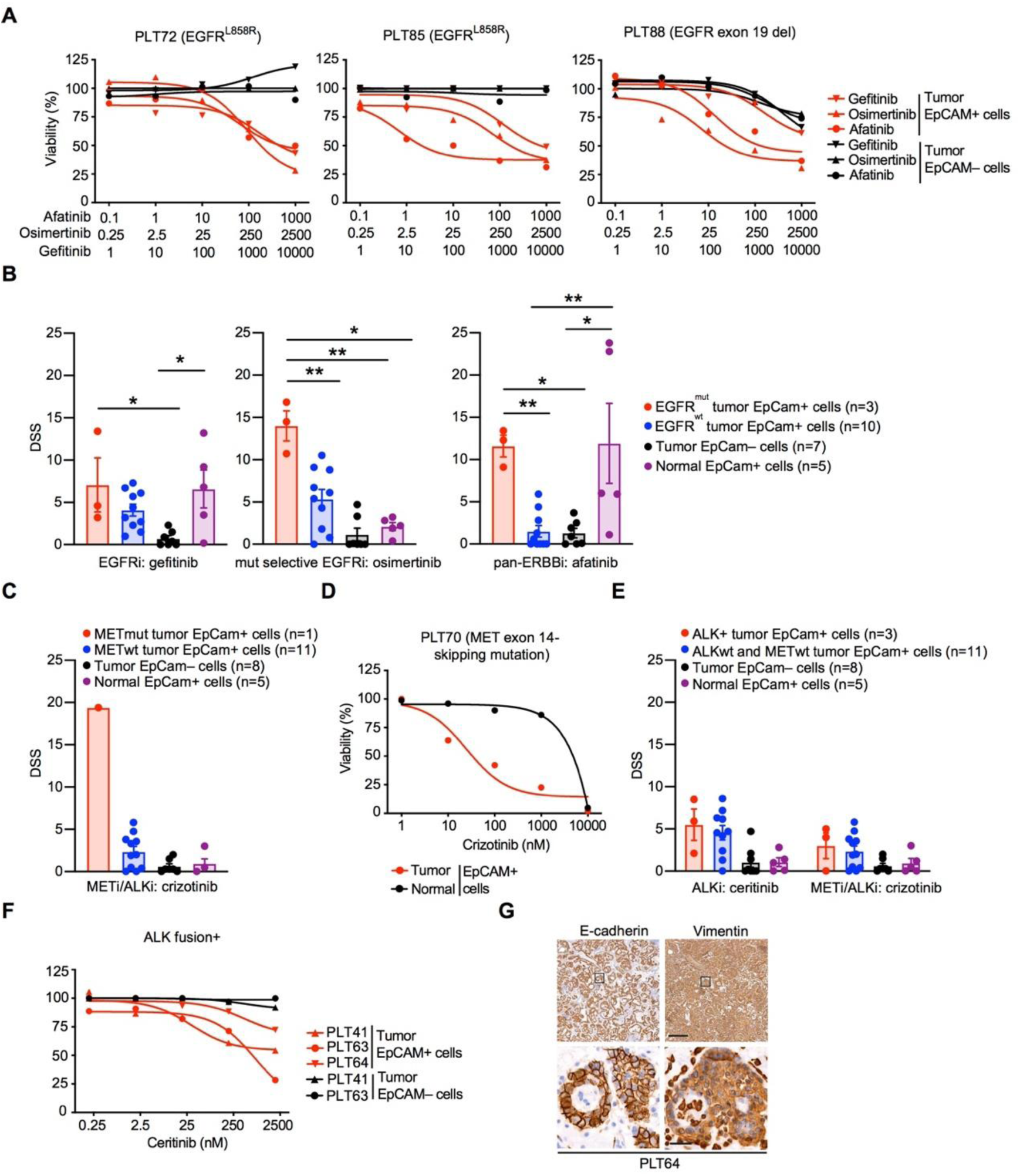
NSCLC FUTCs predict genotype-matched therapeutic responses. (A) Dose-response curves of gefitinib, osimertinib, and afatinib in patient-matched tumor-derived EpCAM+ and EpCAM– cells. (B) DSS for gefitinib, osimertinib, and afatinib compared between *EGFR* mutant EpCAM+, *EGFR* wildtype EpCAM+, tumor EpCAM– and normal EpCAM+ cells. Each dot represents an independent sample. (C) Bar graph representing the DSS of crizotinib and (D) ceritinib and crizotinib across all the patient samples. (E) Dose-response curves of crizotinib in patient-matched tumor-derived EpCAM+ or normal tissue-derived EpCAM+ cells. (F) Dose-response curves of ceritinib in patient-matched tumor-derived EpCAM+ and EpCAM– cells. (G) Representative IHC images of E-cadherin and vimentin staining in patient (PLT64)-derived EML4-ALK+ lung tumor tissues. Scale bars correspond to 200 µm or 20 µm for low or high magnifications, respectively. Boxes indicate areas shown in the higher magnification in a lower row. Error bars represent ± SEM. Two-tailed unpaired Student’s *t* test *p* values are *p<0.05, **p<0.01, ***p<0.001.

Contrasting with the above target-matched responses, EpCAM+ cells derived from three ALK fusion+ tumors showed low sensitivity to the ALK inhibitors ceritinib and crizotinib (Fig 4E-F). Since epithelial-to-mesenchymal transition (EMT) can explain resistance to ALK inhibitors, ^26, 27^ primary tumor tissues were stained with epithelial E-cadherin and mesenchymal vimentin markers (Table S2; six of 18 patient samples showed vimentin staining positive tumor cells). This confirmed EMT in all three ALK+ samples tested in the FUTC assay (Fig 4G and S2E), possibly explaining why these samples demonstrated poor response to ALK inhibition. Thus, FUTC-based profiling can gauge targeted kinase inhibitor responses and non-responses in patient samples that carry the respective driver mutations.

### Pharmacological profiling of *KRAS* mutant patient-derived FUTCs demonstrates inter- and intra-tumoral heterogeneity

We next asked how FUTC-based profiling captures sample-selective functional heterogeneity in *KRAS* mutant samples, which overall represent 25-30% of NSCLC, yet stratify further due differences in, among others, histopathology, metabolism, or co-occurring mutations. Comparison of drug responses of seven *KRAS* mutant and 11 *KRAS* wildtype samples identified the MEK inhibitor trametinib as the most *KRAS* mutant-selective compound, but also other MAPK inhibitors showed KRAS-selective responses, albeit with a less stringent statistical difference (Fig 5A and Table S3). This aligned with analysis of lung cancer cell lines in the GDSC database, ^28^ revealing MAPK inhibitors as the most *KRAS* mutant-selective (Fig. S3A). Moreover, trametinib selectively inhibited *KRAS* mutant cells without affecting tumor-derived EpCAM– and normal tissue-derived EpCAM+ control cells (Fig 5B). Correlation analysis of the drug sensitivities of individual *KRAS* mutant samples to average drug sensitivities of *KRAS* wildtype samples indicated inter-tumoral heterogeneity: each *KRAS* mutant sample exhibited a unique drug response profile, and even for shared hits, such as trametinib, the actual efficacies varied between samples. As anticipated, out of a total of 27 hits, 14 compounds selective for *KRAS* mutant FUTCs either targeted the MAPK or the PI3K-AKT pathways (Fig 5C and S3B).

**Figure 5.**
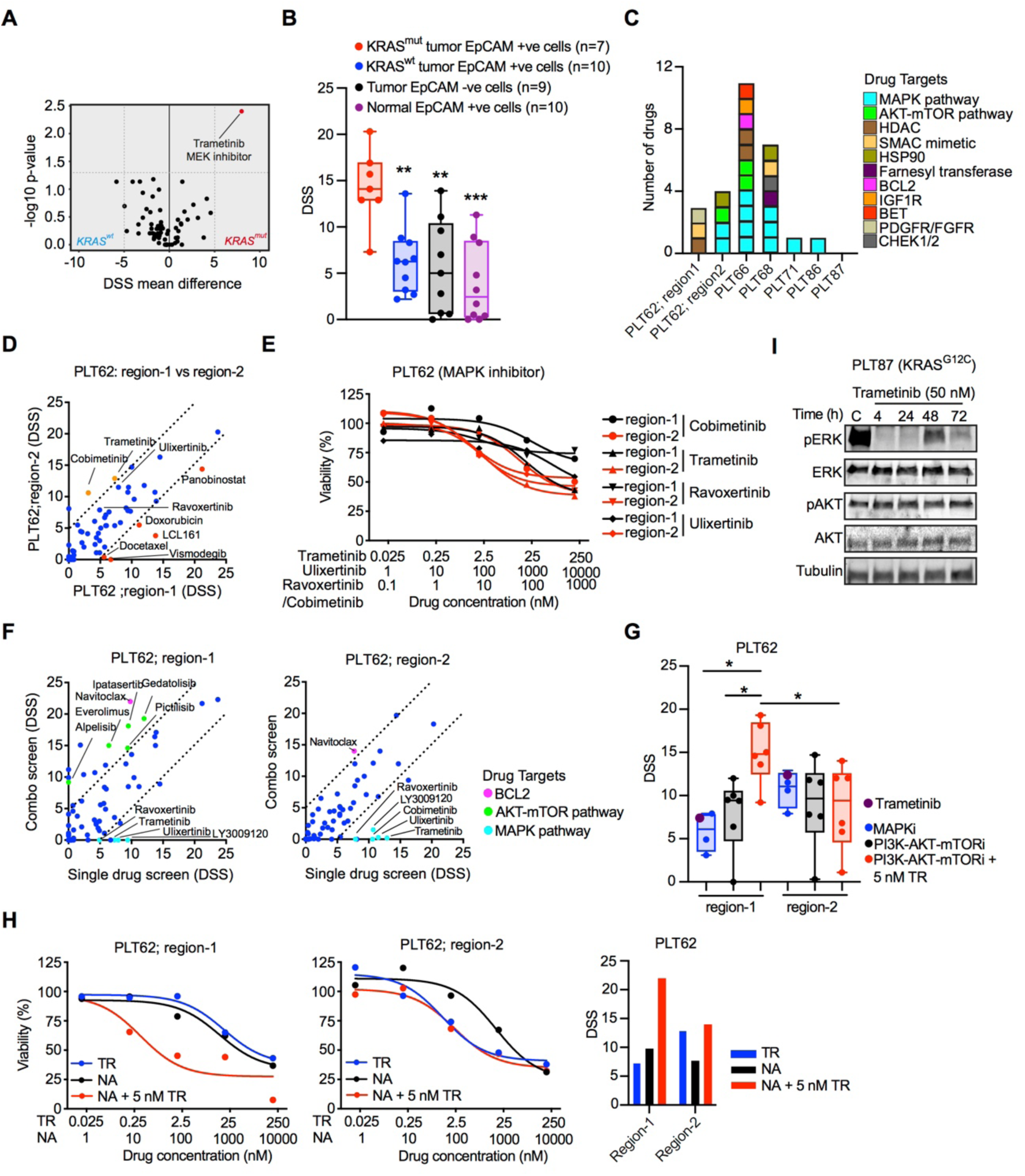
Functional profiling of *KRAS* mutant FUTCs reveals inter- and intra-tumoral heterogeneity. (A) Volcano plot displaying an association between responses to drug candidates and *KRAS* mutation status: DSS comparisons were made between *KRAS* mutant cells (n=7) and *KRAS* wildtype cells (n=11) for 66 oncology drugs using a two-sided Wilcoxon signed-rank test; DSS difference > ± 5 and *p* value < 0.05 were considered as a hit and highlighted with red color. (B) DSS for trametinib (TR) compared between different cell populations; each dot represents an independent sample. (C) The number and target of unique hits identified for an individual patient sample. Boxes are color-coded based on target, and each box represents a single drug. (D) Correlation plots of comparing DSS obtained from FUTCs derived from different tumor regions of the same patient specimen (PLT62). Drug candidates with DSS difference > ± 5 are color-coded with red and orange representing region-1 and region-2 selective hits, respectively. (E) Dose-response curves following indicated MAPK inhibitor treatments in different regions of PLT62. (F) Correlation plots comparing DSS of 66 oncology drugs as a single agent or in combination with 5 nM trametinib. (G) Dot plot displaying DSS of single treatments of MAPK inhibitors (n=4) and PI3K-AKT-mTOR inhibitors (n=6) as a single agent or in combination with 5 nM trametinib. (H) Dose-response curves and DSS bar graph of trametinib, navitoclax (NA), and combination treatment. For the combination screen, 5 nM of trametinib was used together with the dose series of NA. (I) Immunoblots of PLT87-derived *KRAS* mutant FUTCs treated with vehicle (C; DMSO) and or treated with 50 nM trametinib for various time points (4, 24, 48, and 72 h), and probed with indicated antibodies. Error bars represent ± SEM. Two-tailed unpaired Student’s *t* test *p* values are *p<0.05, **p<0.01, ***p<0.001.

To assess intra-tumoral functional heterogeneity, drug responses in FUTCs derived from two different regions from the same patient sample (PLT62) were compared. Both tumor regions carried identical genetic alterations (Fig S4A), including *STK11* and *NFE2L2* mutations known to confer resistance to MEK inhibition in *KRAS* mutant lung cancer. ^21, 29^ A combinatorial screen of 66 drugs in combination with trametinib showed that region-2 cells exhibited a relatively higher sensitivity to MAPK inhibitors (Fig. 5D-E), while region-1 cells showed selective sensitivity to combinatorial treatment with trametinib plus PI3K-AKT pathway inhibitors (Fig. 5F-G). Consistent with the drug sensitivity data, cancer cells from region-1 exhibited a higher percentage of pERK and p4EBP1 positive cells, as well as cells dually stained for these markers, jointly indicating a potential tumor region-selective co-dependency on MAPK and PI3K-AKT pathways (Fig S4B-C). Interestingly, FUTCs from both tumor regions showed combinatorial response to trametinib plus the BCL-2/BCL-xL inhibitor navitoclax (Fig. 5H), a combination described to convey synthetic lethality in *KRAS* mutant tumors. ^30^

The above shows that FUTC profiling can expose inter- and intra-tumoral functional heterogeneity in *KRAS* mutant NSCLC (Fig. 5C-G and Table S2), and tentatively indicates their use in identifying tissue-selective drug combinations. As in murine samples, feedback activation of the MAPK and PI3K-AKT pathways, measured by re-bound ERK and AKT phosphorylation, was detected following trametinib treatment in KRAS mutant FUTCs from PLT87 (Fig 5I). Although analysis of more samples is warranted, this preliminarily results suggests that deeper functional profiling of FUTCs may further help to identify drug combinations that target adaptive signaling mechanisms.

### Compassionate implementation of a FUTC-based functional diagnostic assay

Finally, we report a case where FUTC-based drug response assessments were used to implement compassionate treatment for a patient with refractory metastatic lung cancer. This patient had been diagnosed with *EGFR* mutant stage IV lung adenocarcinoma (AC) 3.5 years earlier and had since been treated with both erlotinib and osimertinib, yet experienced disease recurrence as evidenced by the emergence of multiple nodular lesions in the lungs and widespread metastatic lesions, concomitant with increases in the levels of carcinoembryonic antigen (CEA) and neuron-specific enolase (NSE) tumor markers. At this point, a metastatic lesion from the neck region was surgically resected, and FUTCs were utilized for replicate testing of 233 relevant drugs or drug combinations (Fig. 6A-C).

**Figure 6.**
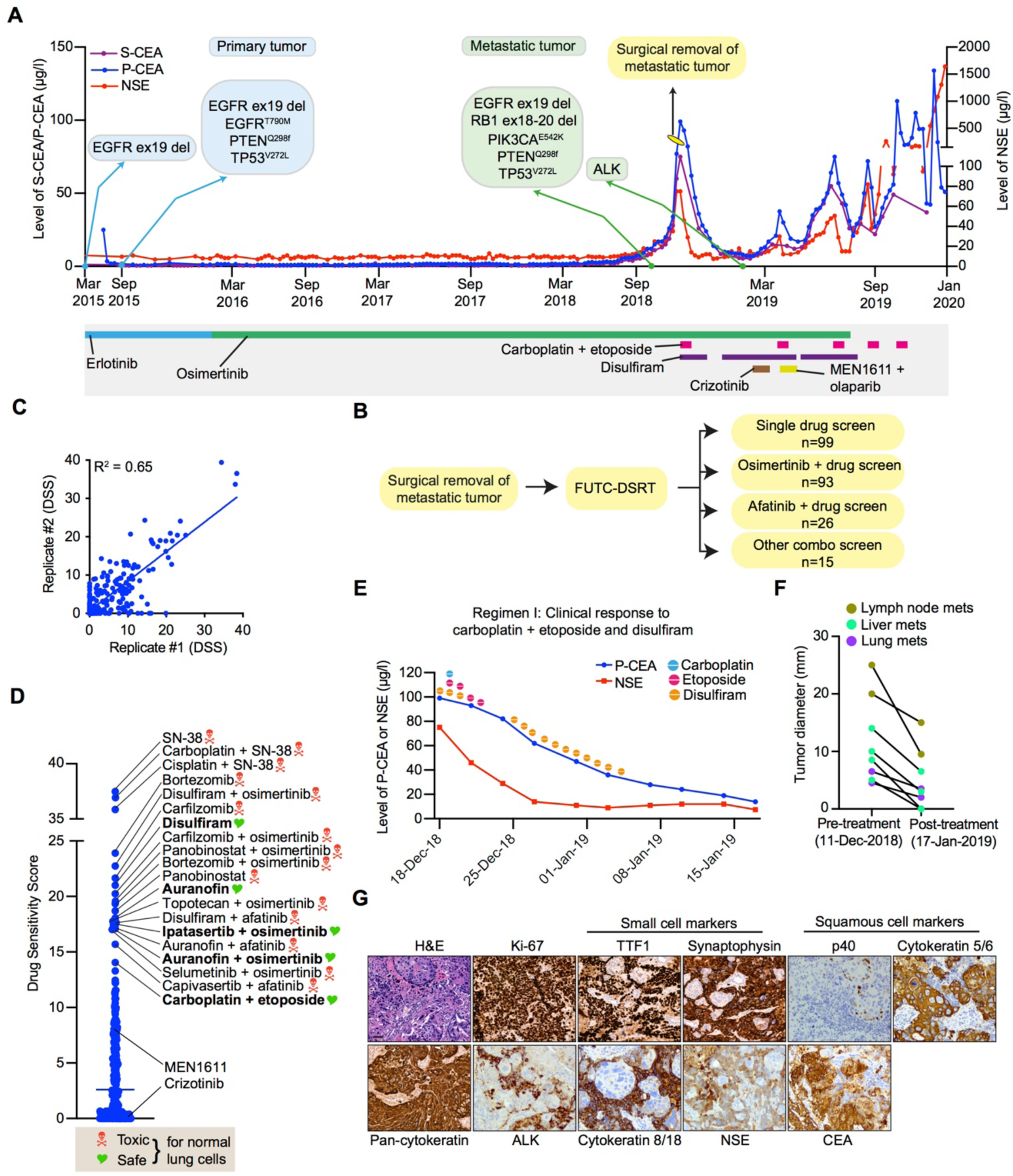
FUTC-based functional testing guided treatment of a patient with refractory lung adenocarcinoma. (A) Treatment outline and measured CSE and NSA levels of a patient with *EGFR* mutant lung adenocarcinoma. (B) A surgically removed tissue specimen was utilized for FUTC-based drug testing of 233 single drugs or drug combinations in duplicate. (C) Correlation plot comparing the DSS from the two replicate tests. (D) The drug sensitivity scores of the top 20 hits with general toxicity based on activity on normal cells noted. (E) Changes in the level of CEA, NSE, and (F) tumor size after carboplatin plus etoposide treatment. (G) Representative images of hematoxylin and eosin (H&E) and IHC staining performed on surgically resected tumor tissue.

The drug sensitivity screen demonstrated poor response to multiple EGFR inhibitors (Fig. S5A), in line with the acquired clinical resistance to erlotinib and osimertinib. To identify effective treatment options with minimal generic toxicities, we first analyzed responses of the top 20 treatments (Fig. 6D) in six patient-derived normal lung CR cultures, established from independent patients (Fig. S5B). This allowed narrowing down of the list and eliminated 15 treatments with generic toxicities. From this analysis, we identified five treatments with selective potency towards the tumor-derived cells, including disulfiram as well as carboplatin combined with etoposide (Fig. S5C). Conversely, the FUTC-based profiling results were in agreement with the clinical non-responses to therapies targeting PI3K and ALK, even though the tumor cells harbored activating mutations in *PIK3CA* and showed overexpression of *ALK* (Fig. 6G and S5D-E).

Based on these results, the scheduled carboplatin plus pemetrexed, which showed inferior sensitivity to carboplatin plus etoposide, was changed and the patient received disulfiram and five cycles of carboplatin plus etoposide with or without disulfiram. This resulted in a therapeutic benefit of the FUTC-defined chemotherapy during the first three treatment cycles, detected by a decrease in tumor size analyzed by computed tomography imaging, as well as reduced levels of CEA and NSE (Fig 6E-F and S6A-E). Interestingly, immunohistochemical analysis of the tumor tissue done a few months following treatment onset revealed two spatially intermixed subtypes of tumor populations: a (i) small cell lung cancer (SCLC) population with NSE expression and (ii) a squamous cell carcinoma (SCC) population with CEA expression (Fig 6G). Histological transformation to SCLC or SCC are known acquired resistance mechanisms in *EGFR* mutant AC patients treated with EGFR inhibitors, ^31, 32^ and platinum plus etoposide is an effective treatment strategy for SCLC patients, ^33, 34^ which the FUTC-based drug testing had identified independently of the histological analyses. However, in line with the poor prognosis of SCLC, during the last two treatment cycles of carboplatin plus etoposide, NSE levels showed an initial drop and then a steep rise, while CEA remained stable, indicating eventual chemoresistance and progression of the SCLC subpopulation, confirmed by pathology analysis of a new recurrent metastatic lymph node biopsy (collected Jan 2020; Fig. S6F-G). This compassionate case therefore demonstrates that therapeutic screening of FUTCs can be used to identify effective treatments many of which cannot otherwise be predicted based on molecular diagnostics, in an individualized manner.

## DISCUSSION

The past decade has seen major advances in the area of precision cancer medicine and spearheaded the development of diagnostic innovations, which includes solutions for *ex vivo* propagation of carcinoma cells. However, unlike murine tumor cells which generally adapt well to primary cultures, the majority of patient-derived cultures we established represented non-malignant cells, corroborating conclusions from other recent studies. ^17, 35, 36^ While the success rates of establishing cultures from pleural effusion are reportedly higher, ^4, 13^ their use is limited as only around a tenth of all of NSCLC patients present with malignant pleural effusions. ^37^ Other preclinical models such as tumor organoids or xenografts also have suboptimal success rates of 20-70% or 30-40%, respectively, ^11, 15, 16, 38, 39^ but these have an additional issue of long establishment times of one to three months for organoids or two to ten months for xenografts. ^4, 11, 15, 16, 38^ Prolonged *ex vivo* propagation in artificial conditions may cause a functional drift of cancer cells from their original identity, and also the patient’s tumor may evolve during this time, further compromising the compatibility of preclinical cultures with the diagnostic clinical decision-making process.

To overcome such limitations, and inspired by precision medicine approaches using liquid biopsy-sampled leukemic cells, we set out to assess whether uncultured tumor cell populations could be utilized for pharmacological profiling immediately following isolation, prior to their attachment to cell culture plates - the FUTC approach. We asked whether the study of epithelial cells in solution might be compromised e.g. due to apoptotic priming related to anoikis. Encouragingly, both murine and human FUTCs demonstrated sustained or increased metabolic activity during the first three days of *ex vivo* culture. In addition, murine *Kras* mutant FUTCs were shown to mimic the MEKi-associated resistance bypass mechanisms and related drug sensitivities previously identified in primary cultures derived from these models. These validating findings supported further translational implementation of the approach, and target-matched drug responses were indeed confirmed in FUTCs derived from surgical tissue samples carrying mutant *EGFR*, *MET,* or *RAS*, or rearrangement in *ALK*, with a positive correlation between cancer cell percentage and drug screen data quality. In KRAS-driven tumors, heterogeneity was observed at both inter- and intra-tumoral levels, which could be explained by underlying differences in biologies such as co-occurring mutations, ^21^ metabolic dependencies explained e.g. by *KRAS* allelic copy gains, ^40^ or phenotypes influenced by the tumor progenitor cells, such as tumor histopathology or oncogenic signaling activities. ^10, 18^ These data overall indicate that the FUTC assay holds promise for further diagnostic development, particularly for samples with high cancer cellularity.

In previous studies, *ex vivo* drug response testing on heterogeneous primary cancer and stromal cell mixtures failed to culminate in clinical benefits, possibly because these earlier assays did not involve the enrichment of tumor cells. ^41, 42^ In the FUTC assay, cancer cell-selective drug profiling is rather done on EpCAM+ epithelial cells isolated from tumor tissue, with parallel analysis of epithelial cells from adjacent normal tissue serving as a toxicity indicator. A recent study relatedly demonstrated that dynamic BH3 protein profiling reliably reveals drug-elicited apoptotic pathway signaling in freshly isolated breast and colorectal cancer tumor cells, further underscoring our conclusions. ^43^ A caveat of the FUTC approach is that not all tumor epithelia express EpCAM, with 40-85% of ACs and 85-98% of SCCs reported as EpCAM positive. ^44–47^ Nevertheless, in our dataset the majority of tested samples (eight out of eight) showed enrichment of cancer cells following EpCAM selection, and while alternative purification methods are being investigated, EpCAM remains the gold standard for capture of blood-circulating tumor cells. ^48–50^ Implementing the EpCAM approach, reliable drug response data, often matched to the targeted drivers, was obtained from 18 of 19 cases, highlighting assay robustness. A question for future exploration is how the FUTC approach can be adapted to smaller biopsy samples.

The FUTC assay was implemented to guide the compassionate treatment of a stage-IV *EGFR* mutant lung AC patient, with cells derived from an operable metastatic tumor in the neck area. While on EGFR inhibitor treatment, the patient had developed an aggressive disease with multiple metastatic lesions and genomic alterations in among others *TP53*, *RB1*, *PIK3CA*, and *PTEN*, commonly associated with EGFR inhibitor resistance. Even though the combined *TP53* and *RB1* mutations may have suggested a risk for histological transformation to SCLC, ^31, 50^ the official diagnosis was AC at the time of FUTC profiling. Nevertheless, in light of FUTC-based drug sensitivity data and the identified mutations, the patient received carboplatin plus etoposide treatment, generally recommended for treatment of SCLC patients, ^33^ leading to a substantial reduction in tumor burden and level of tumor markers. As far as we know, this represented a rare clinical case, with only one other report describing dual transformation of *EGFR* mutant lung ACs to NSE-marked SCLC and CEA-marked SCC. ^51^ These cell populations were found intermixed in the biopsy sample, as if coexisting in symbiosis, perhaps implying that these were derived from a common progenitor cell. In addition to chemotherapy, the patient also received another FUTC-guided treatment, the acetaldehyde dehydrogenase (ALDH2) inhibitor disulfiram, but it’s dosing was reduced as it associated with delirium, possibly because the patient carried a variant ALDH2*2 allele. ^52^ The *ex vivo* approach also affirmed the previously known resistance to EGFR inhibitors and it prospectively predicted the non-responses to PI3K and ALK inhibitors, for which the patient was later treated based on molecular features. FUTC-based drug testing therefore both predicted the cancer’s intrinsic resistance and sensitivity to clinically actionable regimens.

Collectively, this work demonstrates that tumor-derived FUTCs have promise for clinical application. The fast throughput nature of the assay, bypass of *ex vivo* culture, and ability to guide personalized treatment choices render the assay attractive for wider diagnostic use. However, to more broadly assess the clinical feasibility, more clinical profiling cases would need to be actioned, particularly for tumors that lack targetable drivers or metastatic-stage tumors that lack treatment options. Importantly, since FUTCs allow for the identification of effective combinatorial treatments, the assay can possibly be used to assess polytherapy options to circumvent emergence of resistance to single treatments. Indeed, aligning with our findings, a phase 1 clinical trial to test the safety of a triple combination of osimertinib, platinum, and etoposide to circumvent small cell conversion is underway for *EGFR* mutant patients (NCT03567642). In conclusion, FUTC-based functional profiling enables rapid assessment of tumor-selective drug responses, and shows promise for further development as a personalized diagnostic assay to complement genomic and histopathological profiling.

## MATERIALS AND METHODS

### Preparation of murine lung-tumor derived FUTCs

Experiments involving KP and KL genetically engineered mouse models were conducted following the guidelines from the Finnish National Board of Animal Experimentation (permit number ESAVI/9752/04.10.07/2015) and all procedures were performed as described previously. ^18^ For FUTC isolation, individual lung tumors were manually minced using a sterile scalpel and then enzymatically digested in HBSS containing 2 mg/ml collagenase, 0.3 mg/ml dispase and 10 mg/ml bovine serum albumin (BSA), using a Miltenyi gentleMACS dissociator, as described. ^19^ To separate EpCAM+ cells from tumor tissue dissociates, EpCAM (CD326) MicroBeads-dependent enrichment was implemented (130-105-958, MACS), as per the manufacturer’s instructions. These EpCAM+ FUTCs were then either directly utilized for drug response assessment or for establishment of CR cultures. Detailed CR culture procedures are provided in the Supplementary Method section.

### Drug Sensitivity and Resistance Testing (DSRT) assay for murine NSCLC models

Murine lung tumor-derived FUTCs or CR cultures at passage 4 were used for DSRT following published procedures, ^19^ with minor modifications. Briefly, 2500 FUTCs/well or 1500 CR cells/well were seeded in 384-well plates and were exposed to drugs on the following day. Following 72 h cultivation at 37°C, CellTiter-Glo reagent (Promega) was added, and cell viability was measured using a PheraStar FS plate reader (BMG Labtech). Detailed procedures for drug response testing and data analysis are provided in the Supplemental Information section.

### Patient sample processing

Clinical samples used in this study were collected with patient’s informed consent at the Helsinki University Central Hospital (HUCH) and procedures were conducted in accordance with protocol approved by the Coordinating Ethics Committee of the University of Helsinki (License numbers: 85/13/03/00/2015 and HUS-1204-2019). Within 30 mins after lobectomy, tumor and tumor-adjacent healthy lung tissue samples were transported from the hospital to the cell culture facility in cold HBSS. The tissue sample was divided into four parts, one each for DNA isolation, lysate preparation for western blot, histological analysis, and FUTC isolation. Subsequently, for single cell suspension, tumor tissue was manually minced using a sterile scalpel, and then enzymatically digested using a Tumor Dissociation Kit (Miltenyi, 130-095-929) and a gentleMACS Dissociator (130-093-235) following the manufacturer’s instructions. Single cell suspensions were then subjected to separation of EpCAM+ and EpCAM– cell fractions using Miltenyi’s human EpCAM (CD326) MicroBeads (130-061-101). EpCAM+/– cells derived from tumor tissue, and EpCAM+ cells derived from normal tissue were used for performing DSRT.

### Drug Sensitivity and Resistance Testing (DSRT) of patient-derived cells

Tumor and normal tissue-derived cells (2500 cells per well) were seeded to pre-drugged DSRT plates. Following 72 h incubation at 37°C, CellTiter-Glo reagent (Promega) was added in each well, and cell viability was measured using a PheraStar FS plate reader (BMG Labtech). When the cell numbers were not sufficient for performing the 66-drug screen, drug screening for molecular target-selective drugs (5 - 10) were performed by seeding 2500 FUTCs/well in 384- well plates, and on the following day cells were exposed to drugs. Detailed DSRT procedures are provided in the Supplemental Information section.

### Statistical analyses

To assess statistical significance, a Student’s t test, nonparametric Mann-Whitney test, or a two-sided Wilcoxon signed-rank test (Fig. 6A) were used. The results were considered statistically significant if a p-value <0.05 was observed. Error bars indicate standard deviation or standard error of the mean. Pearson’s correlation coefficients were used to assess the significance of correlations and displayed in the XY plots. All the graphs presented here, including dose-response curve fits were generated using GraphPad Prism 8 (GraphPad Software Inc).

## ACKNOWLEDGEMENTS

We dedicate this work to the memory of Dyanne Søraas, whose courage and willpower continues to inspire us to advance lung cancer diagnostics and treatment. We acknowledge all patients who supported our research by consenting access to clinical specimens. We are thankful to Matti Kankainen and Soili Kytölä for advice on analysis of NGS and ddPCR data, Merja Räsänen for processing patient consents, Ashwini Yadav for assistance on *KRAS* mutation-drug response association analysis, Astrid Murumägi for reagents and guidance on the establishment of the CR culture protocol, and Vishal Sinha for assistance on drug response correlation analysis. We thank the Sequencing Core Facility at FIMM, HiLIFE, at the University of Helsinki for NGS analysis, and the Laboratory of Genetics, HUS Diagnostic Center, HUSLAB, at the Helsinki University Hospital for NGS and ddPCR analysis. We thank the thoracic pathologists who helped with the clinical specimen selection, the Laboratory Animal Centre for husbandry support, the FIMM WebMicroscope facility for scanning histological slides, and the FIMM High-Throughput Biomedicine facility for drug screening resources. We thank members of the Verschuren and Wennerberg labs for support and guidance. The study received financial support from the University of Helsinki Integrative Life Science doctoral program (SST); HUSLAB and the Finnish Medical Foundation (MIM); the Academy of Finland (EWV; grants 307111); Novo Nordisk Foundation (KW; Novo Nordisk Foundation Center for Stem Cell Biology, DanStem; grant no NNF17CC0027852); the Sigrid Jusélius Foundation (KW); and a HiPOC 2020 grant from the University of Helsinki (EWV).

## AUTHOR CONTRIBUTIONS

SST, KW, and EWV conceived and designed the study; SST performed experiments for the development of the FUTC-based drug screening assay; JB, AH, and NL performed experiments related to IHC analysis; SST, SP, and JB performed data analyses; LS advised on compassionate case study design; SST, EWV and KW, conducted data interpretation; JR, AK, and MIM assisted in collecting clinical data, received patients’ consent, and the primary tissue workflow; MIM and JL performed clinical pathology analyses; SST, KW, and EWV wrote the manuscript.

## Supplementary Methods

### CR cultures

Both murine and human EpCAM+ cells were propagated using a Conditional Reprogramming (CR) culture protocol. In brief, EpCAM+ cells were plated on irradiated (30 gray) 3T3 cells in F-medium composed of 1:3 v/v DMEM : F-12 nutrient HAM supplemented with 5% FBS, 10 ng/ml EGF (BD Biosciences; 354052), 5 µg/ml insulin (Sigma; I2643), 24 µg/ml adenine (Sigma; A2786), 0.4 µg/ml hydrocortisone (Sigma; H4001), 10 ng/ml cholera toxin (List Biological laboratory; 100B), and 10 µM ROCK inhibitor (Y-27632; ENZO). When CR cultures reached 80% confluence, they were differentially trypsinized using a two-step procedure the first to remove loosely attached feeder cells, and the second to trypsinize epithelial cells. To culture 3T3 cells, we used DMEM supplemented with 10% FBS.

### Drug Sensitivity and Resistance Testing (DSRT) assay

**(A) Murine samples:** Tumor-derived fresh uncultured EpCAM+ cells and CR cultures at passage 4 were used for performing DSRT, as described previously. ^1^ Briefly, 2500 FUTCs or 1500 CR cells per well were seeded in 384-well plates using a Biotek MultiFlo FX RAD dispenser in 20 µl F-medium. Following 24 h incubation, drugs were manually dispensed in 10 µl F-medium at eight concentrations, covering a 10,000-fold concentration range in triplicates. Additionally, multiple replicates of cells were treated with 0.1% dimethyl sulfoxide (DMSO) or 100 µM benzethonium chloride, serving as negative and positive controls, respectively. Following 72 h incubation at 37°C, 30 µl CellTiter-Glo reagent (Promega) was added in each well, and cell viability was measured using a PheraStar FS plate reader (BMG Labtech).

**(B) Patient samples:** To screen 66 lung cancer selective drugs, DSRT plates were prepared in advance by dispensing compounds into 384-well plates (3712, Corning) using an Echo 550 liquid handler (Labcyte), at five concentrations covering a 10,000-fold concentration range. As negative and positive controls, 0.1% DMSO and 100 µM benzethonium chloride were included in wells scattered across the plates. Pressurized Storage Pods (Roylan Developments Ltd.) were used to store pre-drugged DSRT plates and were used within one month. Tumor- and normal tissue-derived cells (2500 cells per well) were seeded in pre-drugged DSRT plates using a Biotek MultiFlo FX RAD dispenser, in 25 µl F-medium. Following 72 h incubation at 37 °C, 25 µl CellTiter-Glo reagent (Promega) was added in each well, and cell viability was measured using a PheraStar FS plate reader (BMG Labtech). If insufficient cells for 66-drug screens were recovered, drug screening for molecular target-selective drugs (5 - 10) were performed.

For the compassionate care case study, patient-derived tumor cells were directly utilized for DSRT without performing EpCAM-based immunomagnetic separation step as cytological and histological assessment of frozen tumor tissue sections prior to drug testing initiation revealed high (<90%) cancer cellularity. In brief, tumor tissue within 60 minutes of surgical excision was embedded in Optimal Cutting Temperature (OCT) media and cryosections cut from the OCT block were stained with H&E. Stained sections were analyzed by a clinical pulmonary pathologist.

### DSRT data analysis

To determine screen-to-screen consistency of DSRT profilings, Z-prime factors were calculated by normalizing the raw luminescence values of drug-treated wells with positive and negative controls. Screens showing a Z-prime factor < 0.2 were not considered for further analysis (one of 19 cases). Dose-response curves were plotted using a Marquardt-Levenberg algorithm via the in-house bioinformatic ‘Breeze’ pipeline. ^2^ Next, dose–response curve parameters were employed to calculate the Drug Sensitivity Score (DSS), as described. ^3^

### Genomic sequencing and data analysis (presented in Table 1)

A DNeasy Blood & Tissue kit (Qiagen) was used to extract genomic DNA from healthy lung and tumor tissue samples and from CR cultures (Table 1). Genomic dsDNA (382-500 ng) was fragmented with a Covaris E220 evolution instrument (Covaris) to a mean fragment size of 200 base pairs (bp). For sample library preparation and enrichment, a KAPA Hyper library preparation kit was used, following the SeqCap EZ HyperCap Workflow User’s Guide Version 1.0 (Roche Nimblegen). In brief, pre-capture amplification was performed using 9 cycles, and captures were performed in multiplexes of 3 to 4 samples using 0.667-1 µg of each library. To identify somatic mutations, targeted next-generation sequencing was performed using the Illumina HiSeq2500 system in HiSeq high output mode using v4 chemistry or HiSeq Rapid run mode using v2 chemistry (Illumina), with the NimbleGen Cancer Panel to capture the exons of 578 cancer-related genes. NimbleGen probes used were 120522_HG19_Onco_R_EZ. Instead of SeqCap HE-Oligos, we used xGen Universal Blocking oligo TS mix (IDT). For post-capture amplification, ten cycles were used in two replicate reactions. The library was quantified for sequencing using the 2100 Bioanalyzer High sensitivity kit. Read length for the paired-end run was 2×101 bp. Pre-processing of short read data was done using the Trimmomatic software and included correction of the sequence data for adapter sequences, bases with low quality, and reads less than 36 bp in length. The BWA-MEM algorithm was then used to map paired-end reads passing the pre-processing onto the human reference genome build 38 (EnsEMBL v82). Finally, variants were called according to the GATK best practice for somatic short variants (version 3.5.1), supplemented with cross-sample contamination and sequencing artifact filtering. In the process, tumor samples (tissue and CR culture) were paired with their patient-matched normal samples and variant calls were filtered against a panel of normals generated from the exome data of 24 healthy unrelated Finnish individuals sequenced in-house earlier.

Annotation for single nucleotide variants and short indels was performed using the Annovar tool against the RefGene database. In brief, variants called from samples were filtered for false-positives by removing variants not passing all GATK filters, residing in intronic and intergenic regions, and causing a synonymous or non-frameshift change as well as variant with a minor allele frequency ≥ 1% in the EPS, 1KG, general ExAC (ExAC), East Asian ExAC (ExAC_EAS), non-Finnish European ExAC (ExAC_NFE), Finnish ExAC (ExAC_FIN) databases, strand odd ratio for single nucleotide variants ≥ 3.00, and strand odd ratio for indels ≥ 11.00, coverage ≤ 10, and variant quality value ≤ 40. Finally, variants were removed if the variant allele frequency between the tumor and normal was < 2%. Tools used in the variant calling process and their versions have been outlined earlier. ^4^

### NGS analysis of lung cancer biopsies in the clinic (presented in Table S1)

Upfront NGS screenings were routinely performed for all the AC and ASC patients but not for SCC patients at Helsinki University Central Hospital (HUCH) before treatment initiation or surgery. DNA was extracted from paraffin samples after pre-treatments with the Maxwell RSC (Promega), according to the manufacturer’s instructions. The concentration of the DNA samples was measured using the NanoDrop ND 1000 (Thermo Fisher Scientific). Amplification-based NGS was performed to identify mutations in all exons of *PIK3CA, EGFR, KIT, KRAS, MET, NRAS,* and *PDGFRA* as well as exons 11-15 of *BRAF*-gene by using an in-house gene panel. Briefly, multiplex PCR was performed with 10 ng of genomic DNA, and then adapters were ligated to each PCR product. The amplicon libraries were constructed, and the quantity of amplicon libraries was determined using the Ion Library TaqMan Quantitation Kit (Thermo Fisher Scientific). Each library was diluted to a concentration of 10 pM, and pooled in equal volumes. Template preparation was performed with an Ion Chef Instrument, and sequencing was carried out on an Ion S5 System with PI Chip. Data was generated using the Torrent Suite Software version 5.8 (Thermo Fisher Scientific). The Ion Reporter Software version 4.6 (Thermo Fisher Scientific) was used to filter out non-coding and polymorphic variants. Mean sequencing depth ≥ 1000 was considered as successful sequencing and called variants were only accepted if allele frequency ≥ 1%. All variants listed after filtering were visualized in the Integrative Genomics Viewer (IGV) to manually discard alterations generated by incorrect calling.

### *KRAS* mutation analysis using digital droplet PCR

Snap-frozen cell pellets of tumor-derived EpCAM+ and EpCAM– cells were utilized to detect *KRAS* mutation variant allele frequencies using digital droplet PCR. DNA was extracted from cell pellets with the Maxwell RSC (Promega), according to the manufacturer’s instructions. The concentration of the DNA samples was measured using a NanoDrop ND 1000 (Thermo Fisher Scientific). Targeted WT and mutation probes for the *KRAS* mutation sites G12 and G13 were designed according to Bio-Rad specifications (www.biorad.com). For each reaction, 2 µl of the extracted DNA was used and performed in duplicate. The QX200 Droplet Generator partitioned the samples (20 µl into 20,000 droplets) for PCR amplification. Following amplification using a thermal cycler, droplets from each sample were analyzed individually on the QX200 Droplet Reader, where positive and negative droplets were counted to provide absolute quantification of the target DNA in digital form. The results were analyzed with the QuantaSoft Analysis Pro Software (v.1.0, Bio-Rad).

### Western blotting

Reference tumor tissues, FUTC pellets, or CR cell pellets were lysed with RIPA buffer supplemented with fresh protease and phosphatase inhibitors (Roche). For protein quantification, BCA Protein Assay was used (G Biosciences; 786-570). Western blotting was performed using 20 µg of protein samples, using precast 4–20% polyacrylamide gel (Bio-Rad, 4561096) for electrophoresis and PVDF membrane (Millipore, IPFL00010) for transfer. Western blot membranes were blocked using Odyssey Blocking Buffer (927-40000) at room temperature for 30 min, probed with primary antibodies at room temperature for 2 h or overnight at 4 °C, and finally probed with IRDye secondary antibodies (LI-COR) diluted 1:10000 in Odyssey Blocking Buffer and scanned using an Odyssey infrared imager (LI-COR).

### Immunohistochemistry (IHC) analyses

Tissue processing and IHC procedures were performed as described previously; ^1^ all samples were processed using the equivalent experimental conditions. The Pannoramic 250 digital slide scanner (3DHISTECH, Budapest, Hungary) was used to acquire whole slide scans of stained tumor sections using a 20x objective, and the Pannoramic Viewer (3DHISTECH Ltd) was used at 1:4 magnification to export TIFF images.

To quantify the percentage of malignant epithelial region or cancer cellularity per tumor specimen, H&E stained whole tissue sections were uploaded to the deep learning-based image analysis cloud service, Aiforia (Fimmic Oy, Helsinki, Finland). Regional training sets were manually created by an imaging expert, by selecting regions-of-interest (206 regions) across all H&E-stained samples as two-layer sets, namely a parental layer for whole tumor tissue annotation and a layer for tumor epithelium-only annotation that was further guided by E-Cadherin staining on neighboring sections. The manually annotated tissue regions were used to train the Aiforia algorithm; multiple training-learning cycles were conducted until the predictions on all samples reached a satisfactory outcome. A total of 22 annotations for tissue regions and 184 annotations for tumor epithelial regions were included in the training. The results were validated by a clinical pulmonary pathologist, via the validation functions of Aiforia, and percentages of tumor epithelial regions per specimen were calculated. To quantify the percentages of proliferating cancer cells, both region- and cell-based annotation tools provided by Aiforia were used. A two-layer training set was created, composed of a parental layer of tumor epithelial regions (based on H&E staining) and a cell-based layer to assign (i) proliferating (Ki-67 positive) and (ii) non-proliferating (Ki-67 negative) tumor cells. Multiple training-learning cycles were performed until the deep-learning prediction reached satisfactory outcome and then percentage of Ki-67 positivity were calculated using total number of proliferating and non-proliferating tumor cells.

To assess MAPK and PI3K-AKT-mTOR signaling pathway activities in tumor regions, images stained with phosphorylated ERK, and 4EBP1 were registered to directly adjacent E-cadherin-stained sections, and the overlap of staining was quantified using a spatial quantification and registration image analysis tool Spa-RQ, as described in ^5^.To assess the co-activation between MAPK and PI3K-AKT-mTOR signaling pathways, images stained with phosphorylated ERK and 4EBP1 were registered to each other and the region of staining overlap was quantified by Spa-RQ. Spa-RQ employed a simple thresholding segmentation algorithm, a consistent threshold for each staining was applied to all the samples. The staining specificity and accuracy of the Spa-RQ was validated by a clinical pulmonary pathologist. To assess EMT, all tumor tissue sections were visually inspected for co-positivity of E-cadherin and vimentin in tumor epithelium-only regions. Tissue sections stained for phosphorylated EGFR, ERBB2, or ERBB3 proteins were visually inspected for positivity in tumor only regions.

### List of antibodies

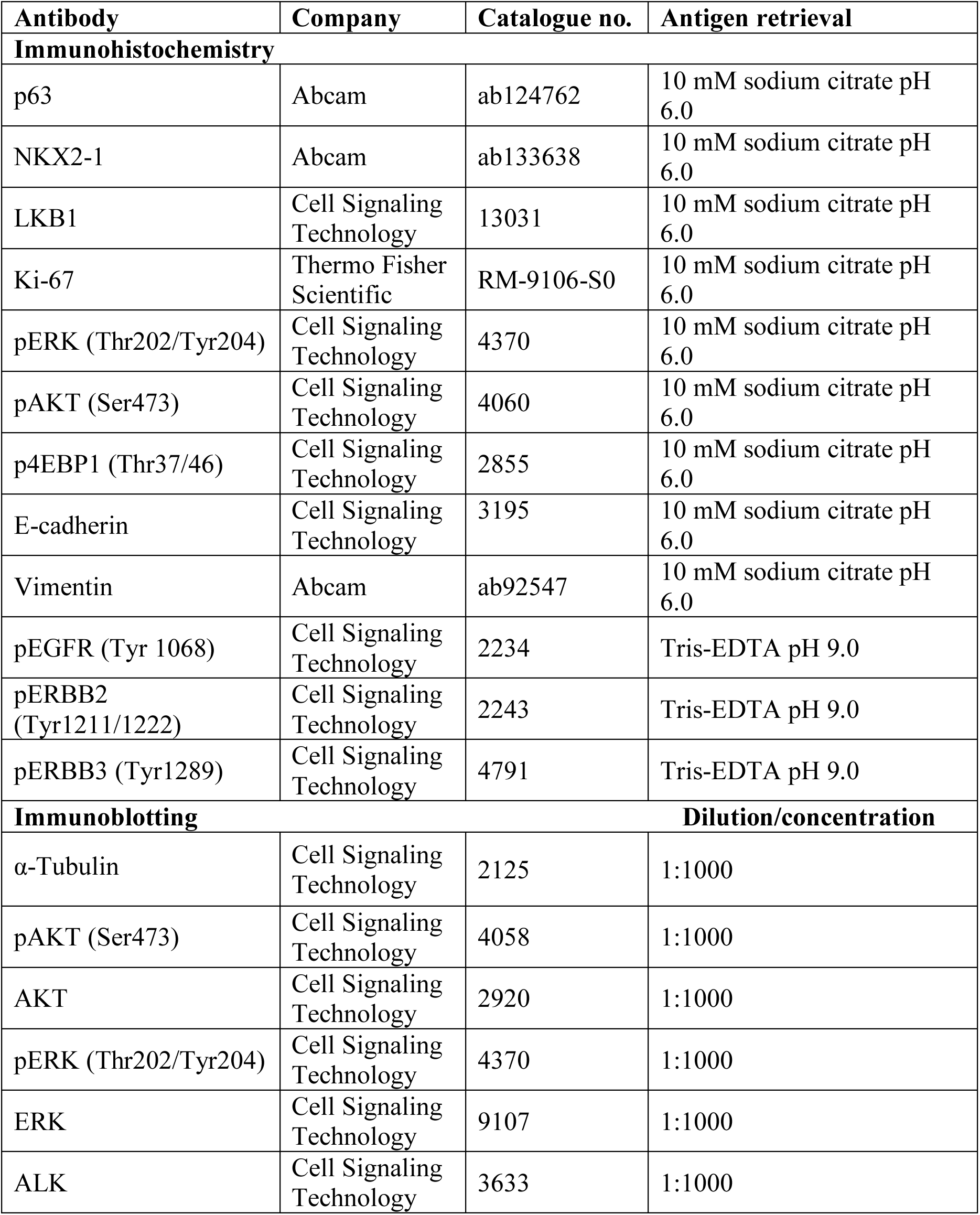

## Supplementary Figures

**Figure S1.**
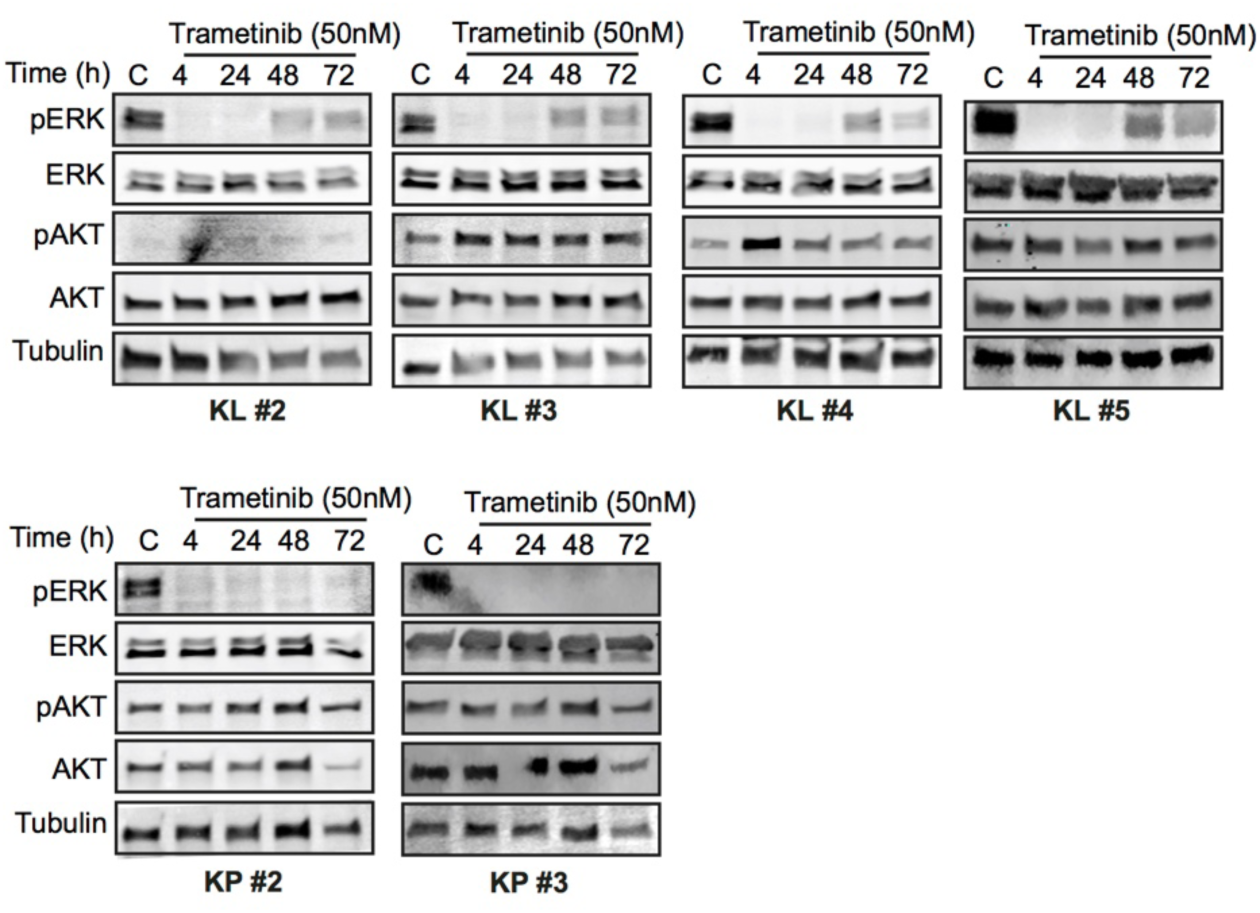
Analysis of trametinib-induced signaling rewiring. Immunoblots of KL and KP FUTCs treated with vehicle (C; DMSO) and or treated with 50 nM TR for various time points (4, 24, 48, and 72 h), and probed with indicated antibodies.

**Figure S2.**
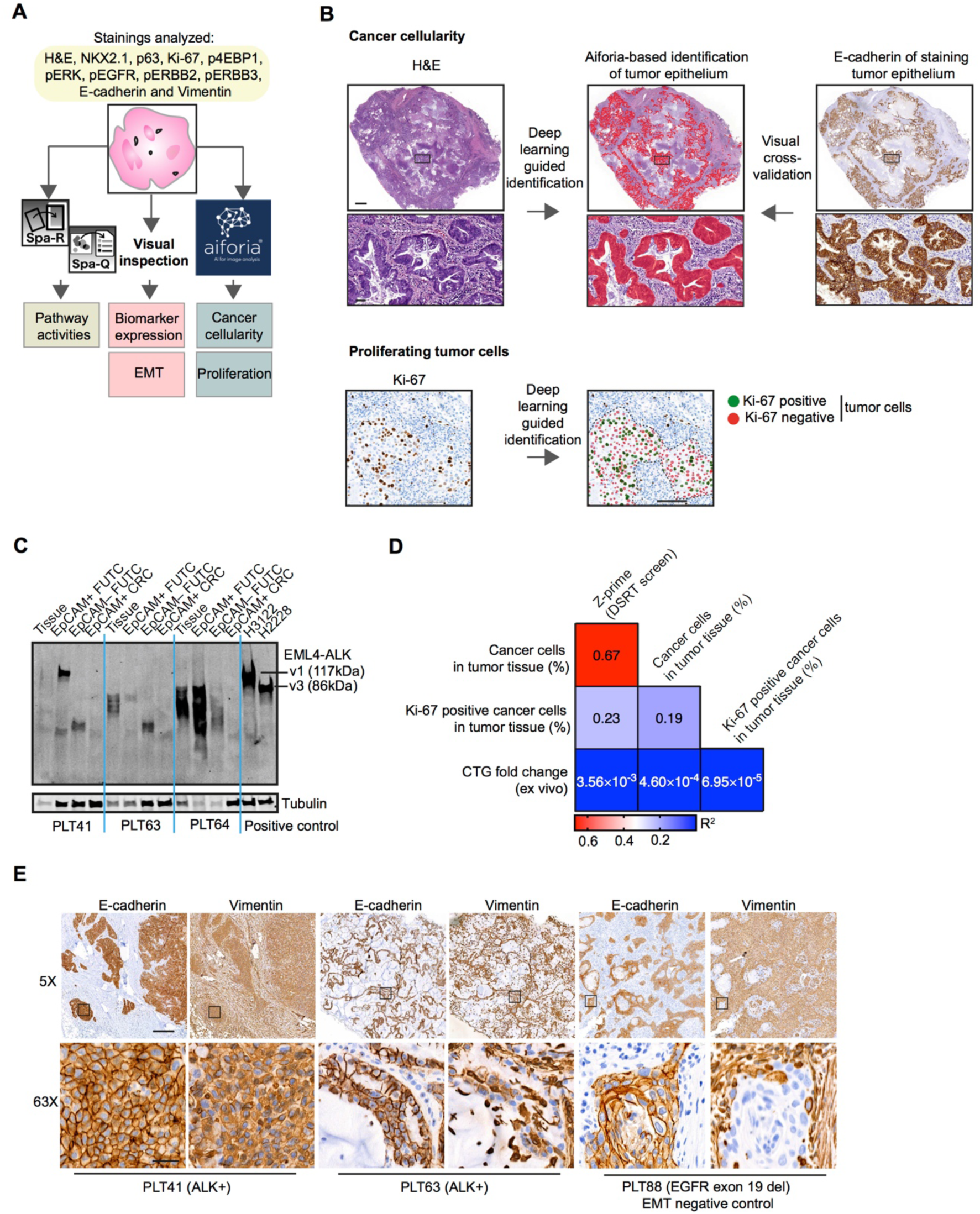
Tumor tissue phenotyping and biochemical analysis of clinical samples. (A) The image analysis software package Spa-RQ^5^ was used for quantification of signaling activities in tumor regions. To identify cancer cellularity and percentages of Ki-67 positive proliferating cancer cells, deep learning-based Aiforia software was used. Biomarker expression and EMT status was visually assessed for each tumor sample. (B) Top panel: representative images demonstrating H&E staining (left), tumor epithelium identified using Aiforia (middle), and the validation via epithelial staining of an adjacent tissue section stained with E-cadherin (right). Scale bars correspond to 1 mm or 50 µm for low or high magnifications, respectively. Bottom panel: representative Ki-67 stained images and Aiforia-based detection of Ki-67 positive (green) and negative (red) tumor cells. The black dotted line indicates the edge of the tumor epithelium region identified by Aiforia in a separate analysis. Scale bars correspond to 100 µm. **(A)** Immunoblot of EML4-ALK+ patient samples analyzed to verify the expression of ALK fusion protein in patient-matched tumor tissue, tumor-derived EpCAM+ and EpCAM– cells, or CR cultures (passage #4). Samples from H3122 and H2228 cells (EML4-ALK+ cell lines) were used as positive controls for v1 (117 kDa) and v3 (86 kDa) variants of EML4-ALK fusion proteins. (D) Heatmap displaying Pearson correlation coefficient values between various factors associated with FUTC-based drug testing. (E) Representative IHC images of E-cadherin and vimentin staining in patient-derived EML4-ALK+ lung tumor tissues. Scale bars correspond to 200 µm or 20 µm for low or high magnifications, respectively. Boxes indicate areas depicted at higher magnification in the bottom row.

**Figure S3.**
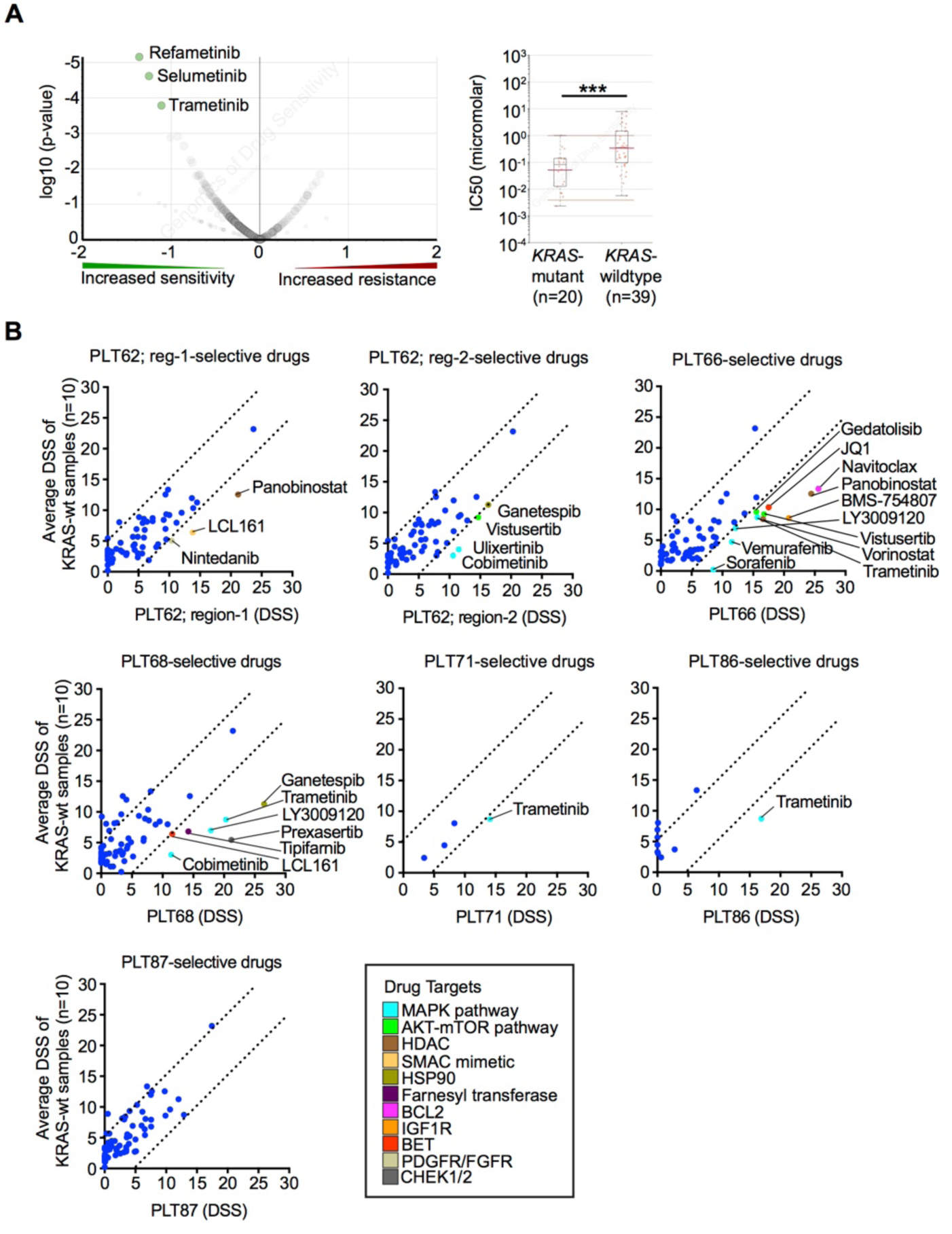
Drug response correlation to identify *KRAS* mutant-selective drug sensitivities. (A) A volcano plot (left) and box plot (right) representation adapted from Genomics of Drug Sensitivity in Cancer (GDSC1) database, showing association between MEK inhibitor sensitivity and KRAS mutation in lung adenocarcinoma cells. (B) Identification of patient-selective drug vulnerabilities by comparison of DSSs of individual *KRAS* mutant samples to average DSSs of *KRAS* wildtype samples (n=10). Drugs showing *KRAS* mutant sample-selective drug sensitivities (DSS difference >5) are color coded based on their target.

**Figure S4.**
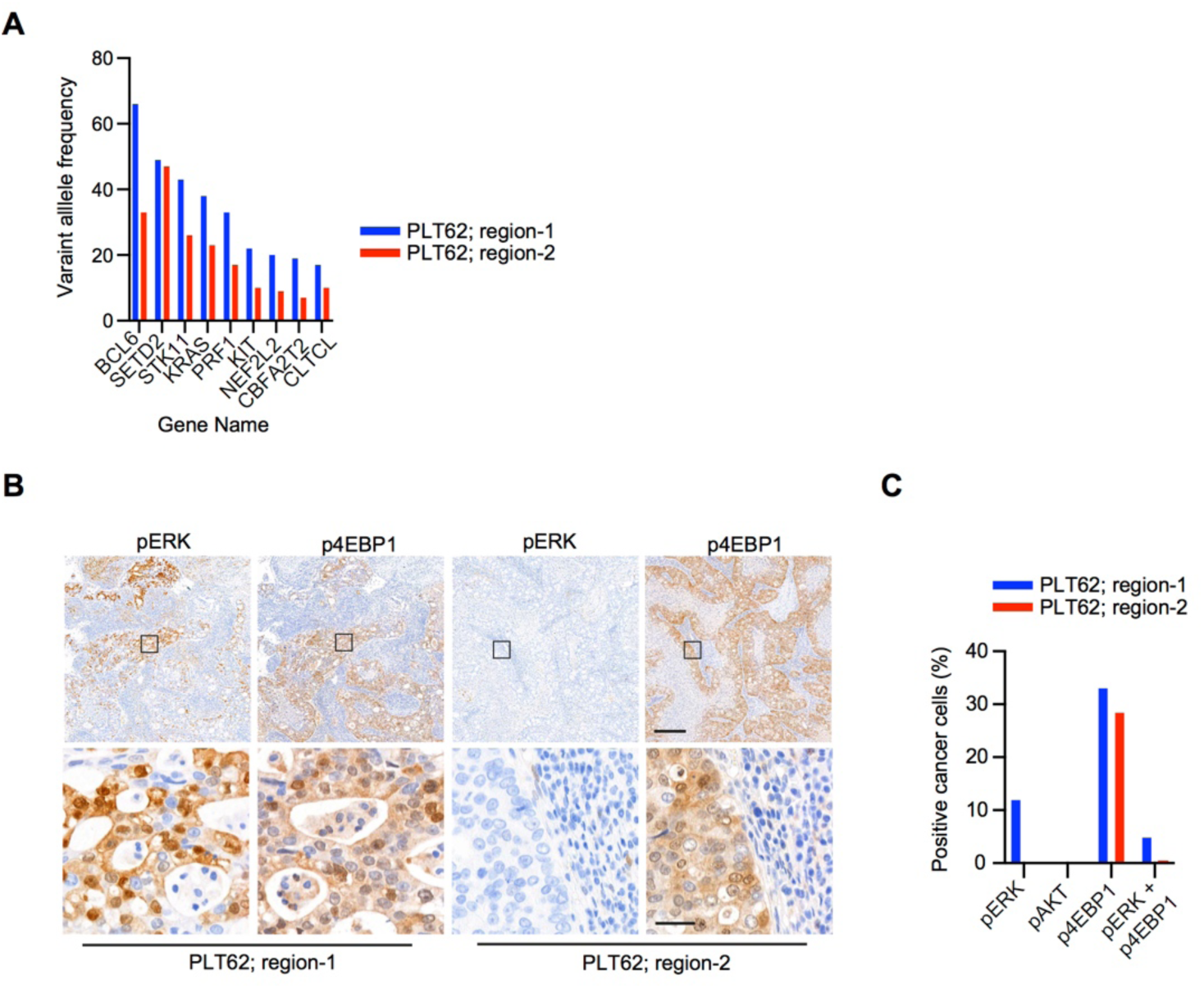
Analysis of intratumoral genomic and phenotypic heterogeneity in a *KRAS* mutant lung cancer specimen. (A) Bar graph displaying the variant allele frequencies of mutations normalized to percentages of cancer cells in their respective tumor region. (B) Representative IHC images of pERK, pAKT, p4EBP1 IHC staining in different regions of the PLT62 tissue sample. Scale bars correspond to 200 µm or 20 µm for low or high magnifications, respectively. (C) Bar graph displaying levels of pERK (Thr202/Tyr204), pAKT (Ser473), p4EBP1 (Thr37/46) and overlapping phosphorylation of ERK and 4EBP1 levels in cancer cells quantified using the IHC image registration and quantification tool, Spa-RQ.

**Figure S5.**
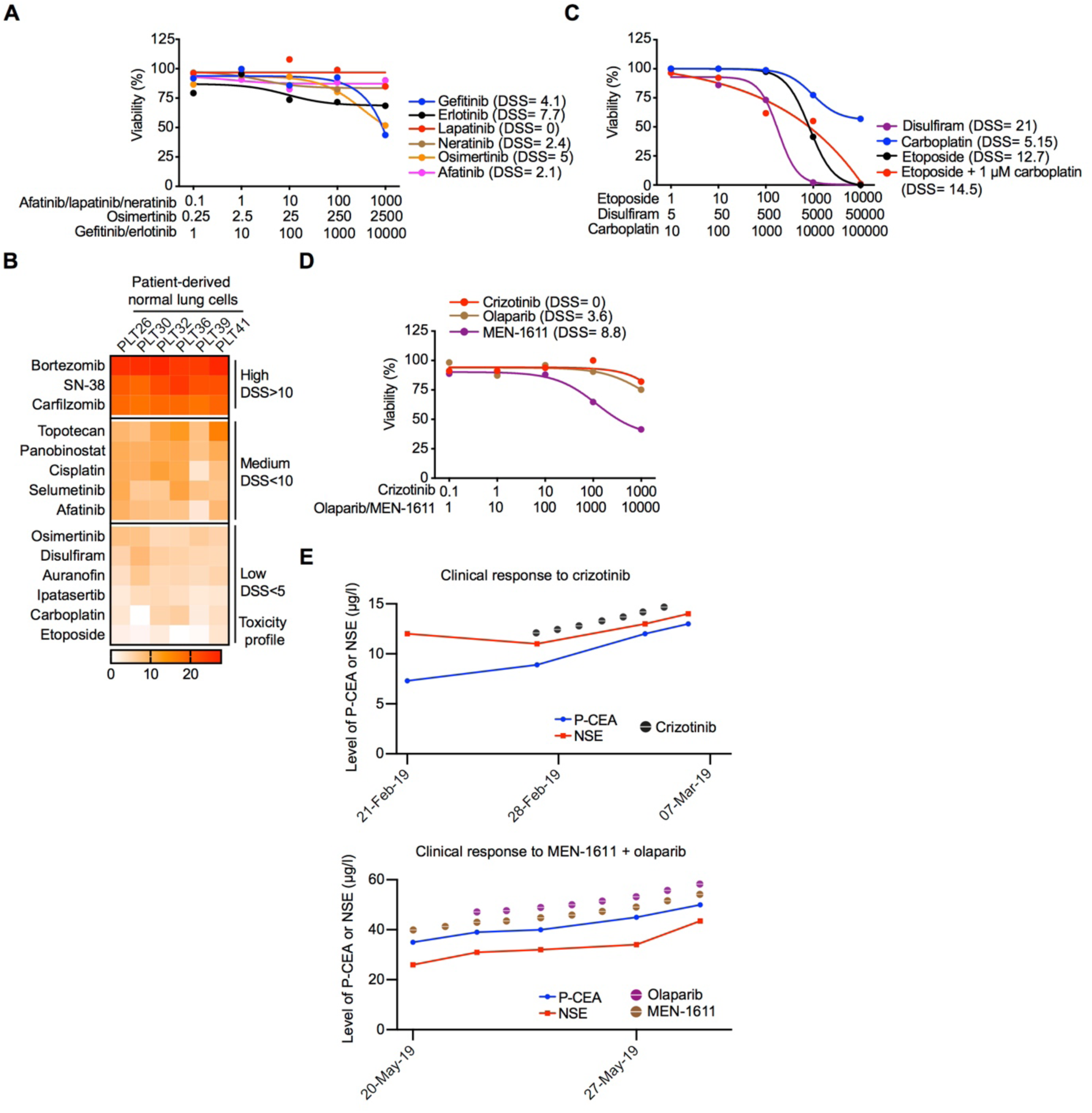
FUTC-based drug profiling identifies patient-selective drug sensitivities. (A) Dose-response curves of different EGFR inhibitors in patient-derived FUTCs. (B) Toxicity profiles of top hits were assessed by testing their sensitivity in CR cultures established from healthy lung tissue of lung cancer patients. Drugs with average DSS >15 or DSS >7 are denoted as highly or mildly toxic, respectively. Drugs with average DSS <7 in the control cells were considered as putative lower toxicity compounds for clinical treatment. (C) Dose-response curves of patient-derived FUTCs treated with five doses of disulfiram, carboplatin, etoposide, and combination treatment. For the combination screen, 1 µM of carboplatin was used together with a dose series of etoposide. (D) Dose-response curves of patient-derived FUTCs treated with five doses of crizotinib, olaparib, and MEN-1611. (E) Changes in the level of CEA and NSE after treatment with crizotinib and MEN-1611 plus olaparib treatments.

**Figure S6.**
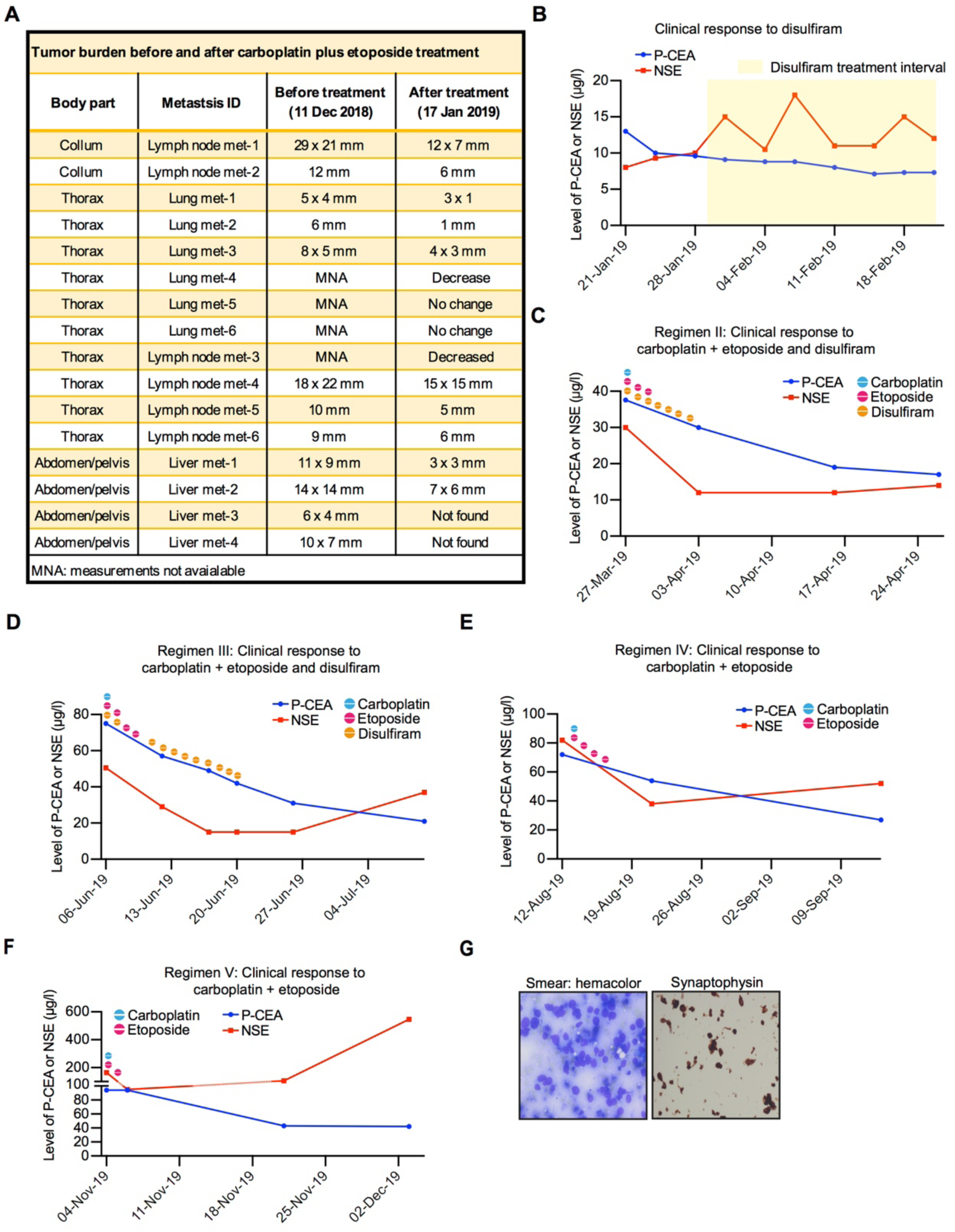
Clinical response to carboplatin plus etoposide treatments. (A) Effect of disulfiram and carboplatin plus etoposide treatment on size of metastatic lesions in lymph nodes, lung, and liver. (B) Changes in the level of CEA and NSE after disulfiram treatments. (C-F) Changes in the level of CEA and NSE after carboplatin plus etoposide treatments with or without disulfiram. (G) Hemacolor stained (left) and synaptophysin (right) stains of cell smear obtained from a fine needle aspiration from a lymph node biopsy (collected in Jan 2020).

**Table S1.** Clinical information of samples used for FUTU-based drug testing

| Sample ID | Sex/age | Lung cancer | TNM stage | Oncogenic driver | Treatment | Smoking history | Tumor location |
| --- | --- | --- | --- | --- | --- | --- | --- |
| PLT37 | F/73 | SCC | T1bN0M0 | nd | surgery | 2 | Left lower lobe |
| PLT38 | M/57 | AC | T3N1M0 | uod | surgery | 0 | Left upper lobe |
| PLT39 | M/66 | MAC | results not available | uod | surgery | 0 | Right upper lobe |
| PLT41 | M/56 | AC | solitary recurr in mediastinum | ALK fusion + | surgery | 0 | Mediastinum |
| PLT62 | F/50 | AC | T2aN1M0 | KRAS G13D | surgery + chemotherapy | 2 | Right lower lobe |
| PLT63 | F/68 | AC | T3N0M0 | ALK fusion + | surgery | 1 | Left upper lobe |
| PLT64 | F/25 | AC | T2aN1M0 | ALK fusion + | surgery + chemotherapy | 0 | Left lower lobe |
| PLT66 | M/71 | AC | T3N0M0 | KRAS G12C | surgery | 2 | Right lower lobe |
| PLT68 | F/68 | AC | T1cN0M0 | KRAS G12S | surgery | 1 | Right lower lobe |
| PLT70 | M/82 | AC | T1cN0M0 | MET exon 14 skipping mutation | surgery | 0 | Right upper lobe |
| PLT71 | F/58 | AC | T2N0M0 | KRAS G12V | surgery | 2 | Right lower lobe |
| PLT72 | M/70 | AC | T2bN0M0 | EGFR L858R | surgery | 1 | Right upper lobe |
| PLT84 | M/60 | AC | T2N2M0 | EGFR exon 19 del | surgery + adjuv. chemotherapy | 0 | Right upper lobe |
| PLT85 | F/76 | AC | T2aN0M0 | EGFR L858R | surgery | 0 | Right upper lobe |
| PLT86 | F/68 | AC | T2N0M0 | KRAS G12V | surgery + adjuv. chemotherapy | 0 | Right upper lobe |
| PLT87 | F/61 | AC | T3N0M0 | KRAS G12C | surgery | 2 | Right lower lobe |
| PLT88 | F/43 | ASC | T2bN1M0 | EGFR exon 19 del and EGFR amp | surgery + postop. chemoradiotherapy | 0 | Right upper lobe |

SCC: squamous cell carcinoma, AC: adenocarcinoma, MAC: mucinous adenocarcinoma, ASC: adenosquamous carcinoma, nd: not done, uod: unknown oncogenic driver

**Table S2.**
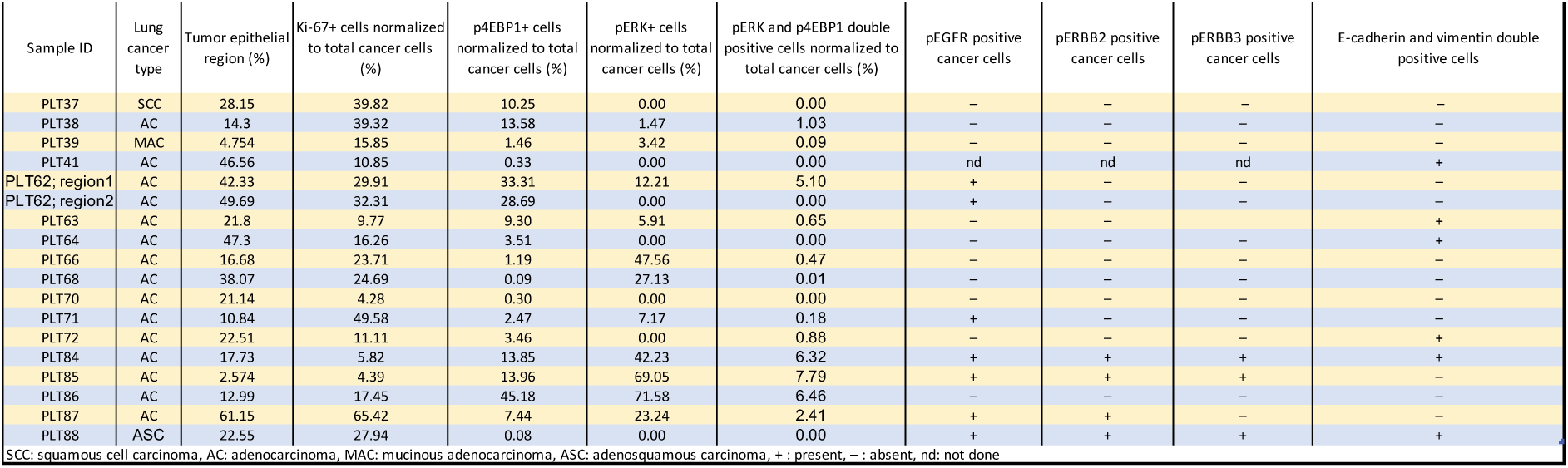
Immunohistochemical and phenotypic characterization of clinical lung tumor tissues

**Table S3.**

| KRAS-selective hit ID | Drug Name | Mean Diff | Median Diff | p.val | adjusted Pval | t.test | t.adjust |
| --- | --- | --- | --- | --- | --- | --- | --- |
| 1 | Trametinib | 8.060 | 7.850 | 0.004 | 0.262 | 0.001 | 0.072 |
| 2 | Panobinostat | 4.633 | 1.350 | 0.268 | 0.937 | 0.245 | 0.737 |
| 3 | LY3009120 | 4.227 | 2.600 | 0.149 | 0.937 | 0.114 | 0.737 |
| 4 | Ganetespib | 3.884 | 2.300 | 0.745 | 0.958 | 0.424 | 0.737 |
| 5 | Cobimetinib | 3.103 | 0.100 | 0.329 | 0.937 | 0.199 | 0.737 |
| 6 | Ulixertinib | 2.573 | 4.400 | 0.191 | 0.937 | 0.370 | 0.737 |
| 7 | Sorafenib | 2.083 | 0.000 | 0.353 | 0.937 | 0.280 | 0.737 |
| 8 | Ravoxertinib | 1.577 | 1.500 | 0.254 | 0.937 | 0.349 | 0.737 |
| 9 | Olaparib | 1.504 | 1.400 | 0.074 | 0.809 | 0.188 | 0.737 |
| 10 | Gandotinib | 1.366 | 3.000 | 0.415 | 0.937 | 0.376 | 0.737 |
| 11 | Prexasertib | 1.340 | 2.850 | 1.000 | 1.000 | 0.771 | 0.950 |
| 12 | Nintedanib | 1.217 | 1.750 | 0.432 | 0.937 | 0.610 | 0.895 |
| 13 | Afatinib | 1.005 | 5.900 | 0.356 | 0.937 | 0.673 | 0.907 |
| 14 | Tipifarnib | 0.996 | 1.150 | 0.745 | 0.958 | 0.652 | 0.906 |
| 15 | LCL161 | 0.963 | 3.600 | 0.569 | 0.937 | 0.801 | 0.950 |
| 16 | Vorinostat | 0.821 | 0.700 | 0.876 | 1.000 | 0.761 | 0.950 |
| 17 | Vismodegib | 0.707 | 0.950 | 0.501 | 0.937 | 0.727 | 0.941 |
| 18 | Gedatolisib | 0.683 | 1.900 | 0.755 | 0.958 | 0.837 | 0.950 |
| 19 | BMS-754807 | 0.669 | 0.050 | 0.935 | 1.000 | 0.848 | 0.950 |
| 20 | JQ1 | 0.363 | -1.300 | 1.000 | 1.000 | 0.914 | 0.955 |
| 21 | Vemurafenib | 0.057 | -1.200 | 1.000 | 1.000 | 0.982 | 0.982 |
| 22 | Dabrafenib | -0.151 | 0.700 | 0.563 | 0.937 | 0.926 | 0.955 |
| 23 | Doxorubicin | -0.244 | 3.000 | 0.639 | 0.937 | 0.946 | 0.961 |
| 24 | AZD4547 | -0.257 | 0.400 | 1.000 | 1.000 | 0.860 | 0.950 |
| 25 | Carboplatin | -0.297 | -0.100 | 1.000 | 1.000 | 0.856 | 0.950 |
| 26 | Dasatinib | -0.314 | -0.050 | 1.000 | 1.000 | 0.896 | 0.955 |
| 27 | Roxadustat | -0.324 | -0.200 | 1.000 | 1.000 | 0.864 | 0.950 |
| 28 | Vistusertib | -0.417 | -2.000 | 1.000 | 1.000 | 0.912 | 0.955 |
| 29 | Chloroquine | -0.500 | -1.550 | 1.000 | 1.000 | 0.808 | 0.950 |
| 30 | PAC-1 | -0.531 | -0.400 | 0.807 | 0.987 | 0.659 | 0.906 |
| 31 | TAK-715 | -0.796 | -2.150 | 0.508 | 0.937 | 0.574 | 0.876 |
| 32 | Sonidegib | -0.881 | 0.000 | 0.440 | 0.937 | 0.315 | 0.737 |
| 33 | Perifosine | -0.884 | -0.750 | 0.456 | 0.937 | 0.296 | 0.737 |
| 34 | Alpelisib | -0.920 | -1.200 | 0.394 | 0.937 | 0.391 | 0.737 |
| 35 | Metformin | -0.923 | -0.500 | 0.238 | 0.937 | 0.183 | 0.737 |
| 36 | Foretinib | -1.003 | -1.450 | 0.608 | 0.937 | 0.447 | 0.737 |
| 37 | Docetaxel | -1.027 | 1.550 | 0.684 | 0.958 | 0.647 | 0.906 |
| 38 | KU-60019 | -1.030 | -0.100 | 0.522 | 0.937 | 0.284 | 0.737 |
| 39 | Volasertib | -1.126 | -1.100 | 0.102 | 0.937 | 0.062 | 0.737 |
| 40 | Cabozantinib | -1.130 | 0.600 | 0.934 | 1.000 | 0.416 | 0.737 |
| 41 | Ipatasertib | -1.176 | -2.600 | 0.755 | 0.958 | 0.704 | 0.929 |
| 42 | Ponatinib | -1.287 | -0.475 | 0.628 | 0.937 | 0.444 | 0.737 |
| 43 | Palbociclib | -1.303 | -1.300 | 0.515 | 0.937 | 0.472 | 0.760 |
| 44 | Veliparib | -1.320 | -0.700 | 0.463 | 0.937 | 0.264 | 0.737 |
| 45 | Osimertinib | -1.428 | -0.950 | 0.518 | 0.937 | 0.446 | 0.737 |
| 46 | Dexamethasone | -1.487 | 0.000 | 0.717 | 0.958 | 0.370 | 0.737 |
| 47 | Birabresib | -1.664 | -2.000 | 0.639 | 0.937 | 0.584 | 0.876 |
| 48 | Gefitinib | -1.677 | 1.300 | 0.792 | 0.987 | 0.389 | 0.737 |
| 49 | Dovitinib | -1.711 | -0.200 | 0.741 | 0.958 | 0.389 | 0.737 |
| 50 | Lapatinib | -1.743 | -0.100 | 0.066 | 0.809 | 0.250 | 0.737 |
| 51 | NVP-LGK974 | -1.813 | -1.550 | 0.623 | 0.937 | 0.217 | 0.737 |
| 52 | Gemcitabine | -1.888 | -2.275 | 0.429 | 0.937 | 0.186 | 0.737 |
| 53 | Crizotinib | -1.902 | -2.700 | 0.136 | 0.937 | 0.069 | 0.737 |
| 54 | Ralimetinib | -1.971 | -1.550 | 0.515 | 0.937 | 0.255 | 0.737 |
| 55 | VE-821 | -2.069 | -1.400 | 0.260 | 0.937 | 0.070 | 0.737 |
| 56 | Ruxolitinib | -2.171 | -1.100 | 0.624 | 0.937 | 0.330 | 0.737 |
| 57 | Ceritinib | -2.492 | -1.725 | 0.228 | 0.937 | 0.201 | 0.737 |
| 58 | Pomalidomide | -2.869 | -2.000 | 0.116 | 0.937 | 0.182 | 0.737 |
| 59 | Vincristine | -2.994 | -1.700 | 0.326 | 0.937 | 0.155 | 0.737 |
| 60 | Azacitidine | -3.344 | -3.650 | 0.073 | 0.809 | 0.069 | 0.737 |
| 61 | Navitoclax | -3.365 | -5.325 | 0.628 | 0.937 | 0.486 | 0.764 |
| 62 | Dinaciclib | -3.503 | -7.100 | 0.432 | 0.937 | 0.438 | 0.737 |
| 63 | Everolimus | -3.991 | -4.000 | 0.074 | 0.809 | 0.061 | 0.737 |
| 64 | Pictilisib | -4.691 | 0.000 | 0.569 | 0.937 | 0.155 | 0.737 |
| 65 | Idasanutlin | -4.938 | -7.300 | 0.202 | 0.937 | 0.082 | 0.737 |
| 66 | Cisplatin | -5.811 | -1.300 | 0.074 | 0.809 | 0.117 | 0.737 |

